# SCRaPL: hierarchical Bayesian modelling of associations in single cell multi-omics data

**DOI:** 10.1101/2021.05.13.443959

**Authors:** Christos Maniatis, Catalina A Vallejos, Guido Sanguinetti

## Abstract

Single-cell multi-omics assays offer unprecedented opportunities to explore gene regulation at cellular level. However, high levels of technical noise and data sparsity frequently lead to a lack of statistical power in correlative analyses, identifying very few, if any, significant associations between different molecular layers. Here we propose SCRaPL, a novel computational tool that increases power by carefully modelling noise in the experimental systems. We show on real and simulated multi-omics single-cell data sets that SCRaPL achieves higher sensitivity and better robustness in identifying correlations, while maintaining a similar level of false positives as standard analyses based on Pearson correlation.

## 1 Introduction

High throughput single cell assays based on next generation sequencing are revolutionising our understanding of biology, with profound implications both fundamental and translational [1]. Single cell technologies avoid the confounding factors emerging from averaging over potentially heterogeneous cell populations [2], providing a global map of biological cell-to-cell variability at the molecular level [3].

While single-cell transcriptomic technologies are rapidly reaching maturity, more recent platforms have emerged that enable simultaneous large scale measurements of multiple molecular layers within the same cell. Multi-omics assays can now capture DNA methylation and gene expression [4, 5], gene expression and copy-number variations [5], and most recently chromatin accessibility along with DNA methylation and gene expression [6] for the same cell. Such platforms have enormous potential to elucidate the mechanisms of gene regulation in unprecedented detail.

Despite the huge potential for breakthroughs, technical limitations in multi-omics technologies create formidable statistical challenges in the interpretations of their results. Single-cell sequencing technologies are notoriously affected by high noise levels, including very strong data sparsity. Such problems are amplified in multi-omics studies, where multiple independent sources of noise might affect the joint distribution of the measurements. Additionally, challenges with normalization strategies, batch effects or other latent variables related to cellular processes might further prevent biological components to emerge clearly from data [7]. As a result, direct adoption of classical statistical tools to assess associations between different molecular layers (e.g. Pearson’s correlation) routinely leads to underpowered analyses, which are only able to identify a handful of significant associations [4, 6, 8].

In this paper, we argue that proper treatment of noise is essential in order to robustly retrieve significant statistical associations. To do so, we introduce SCRaPL (**S**ingle **C**ell **R**egul**a**tory **P**attern **L**earning), a Bayesian hierarchical model to infer associations between different omics components. The Bayesian hierarchical framework, which has already been extensively used in single-omics single-cell analyses (e.g. [9, 10]), explicitly and transparently decomposes noise in the data, enabling efficient extraction of biological signals from technical noise. We demonstrate on both synthetic and real data sets that SCRaPL is both highly accurate and sensitive, identifying much larger numbers of statistically significant associations than standard correlation analyses while retaining a good control on false positives.

## 2 Results

### 2.1 SCRaPL: Single Cell Regulatory Pattern Learning

Single cell multi-omic assays provide unprecedented opportunities to interrogate the relationship between different molecular layers with high resolution. We introduce SCRaPL — a Bayesian hierarchical framework for single cell multi-omics data, inferring cross-layer associations that are robust to technical noise. SCRaPL combines a latent multivariate Gaussian structure with noise models that are tailored to single cell sequencing data (Figure 1). Here, we focus on the links between RNA expression and DNA methylation (DNAm), but additional noise models can be incorporated to accommodate other measurements (e.g. chromatin accessibility).

**Figure 1:**
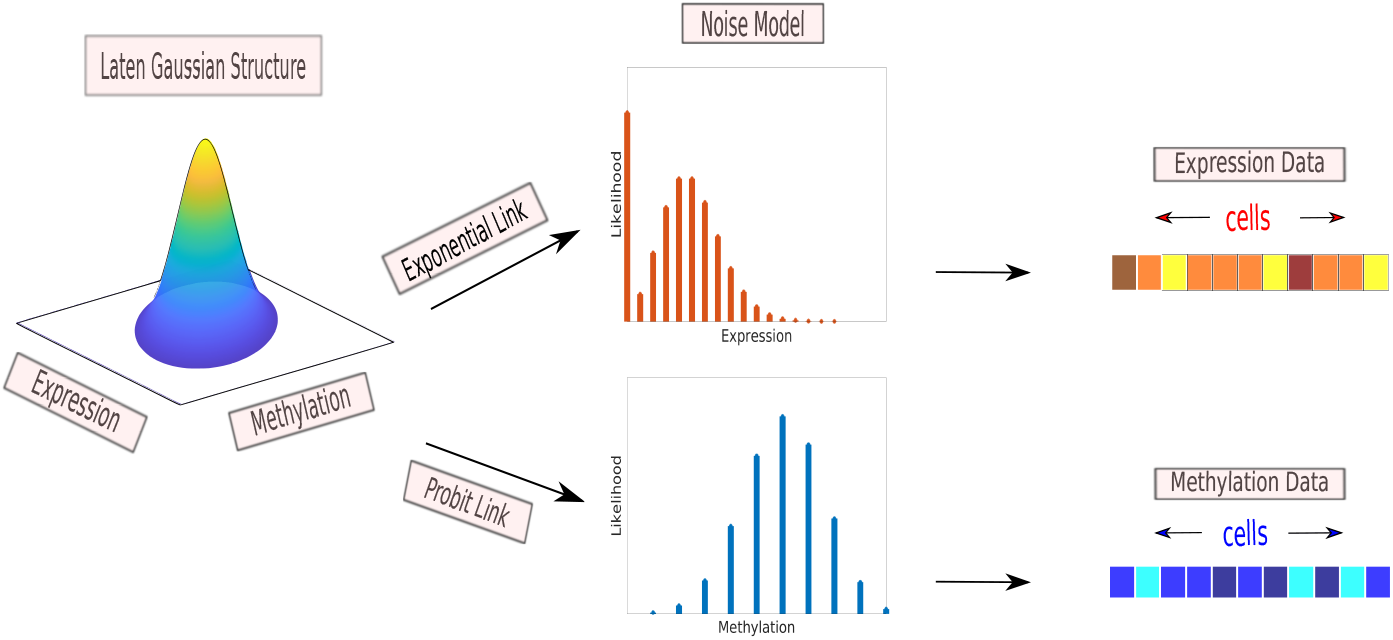
Schematic representation for the SCRaPL model.

For gene expression, Poisson noise is coupled with an exponential link function. This leads to a Poisson-lognormal model, which has been previously used for single cell RNA sequencing (scRNAseq) data [11]. An *over-dispersion* captures variability beyond Poisson noise, leading to the same mean-variance relationship as the negative binomial model (*Methods*). If zero inflation is present [12], a zero inflated (ZI) Poisson noise model can be used [13]. For each gene, evidence of ZI is quantified via the Deviance Information Criteria (DIC, [14]). Moreover, cell-specific scaling factors *s*_*i*_ capture systematic differences across cells. These are inferred using scran [15] and used as fixed offsets.

Matching DNAm data for each gene is quantified as the number of methylated CpGs within a pre-specified genomic region (e.g. gene promoter). To model DNAm data, we use a Binomial noise model which explicitly takes into account the coverage within the region. A probit link function is then used for the latent Gaussian term. Associations between molecular layers are quantified using gene-specific latent correlation parameters *ρ*_*j*_. Statistically significant associations are highlighted using a probabilistic decision rule [16] which controls the expected false discovery rate (EFDR, [17]). More details about this procedure and the algorithm used for parameter inference are provided in the *Methods*. Moreover, a Matlab implementation is available at https://github.com/chrmaniatis/SCRaPL.

### 2.2 Benchmarking SCRaPL using synthetic data

To assess the estimation performance of SCRaPL, we experimented on synthetic datasets covering scenarios with different numbers of cells and a range of values in terms of methylation coverage, ZI for the expression data along with different latent mean and covariance structures (*Methods*, Section 4.8). Here, we focus on estimation accuracy for the gene-specific latent correlation parameters *ρ*_*j*_, but results for other model parameters are displayed in Supplementary Figures S3. We consider a number of simulation scenarios, described in detail in Section 4.8. We start by considering a situation of perfect model specification (experiments 1-3 in Section 4.8), in order to assess the identifiability of our model. In this case, we observe that posterior estimates of correlation tend to be unbiased, with an accuracy which increases with the number of cells in the data set (Figure 2a). As expected, the performance degrades with increasing levels of ZI (Figure 2c). However, we did not observe significant differences across different levels of coverage (Figure 2e). To probe the importance of prior specification, we generated data where the underlying correlation values *ρ*_*j*_ were in an area with low prior mass (experiments 4, 5 and 6 in Section 4.8). In this case, we did observe some bias in our estimates (Supplementary Figure S4), but the latter diminished with increasing sample sizes. Similarly, performance diminishes with increasing ZI levels and stays relatively intact across different coverage levels.

**Figure 2:**
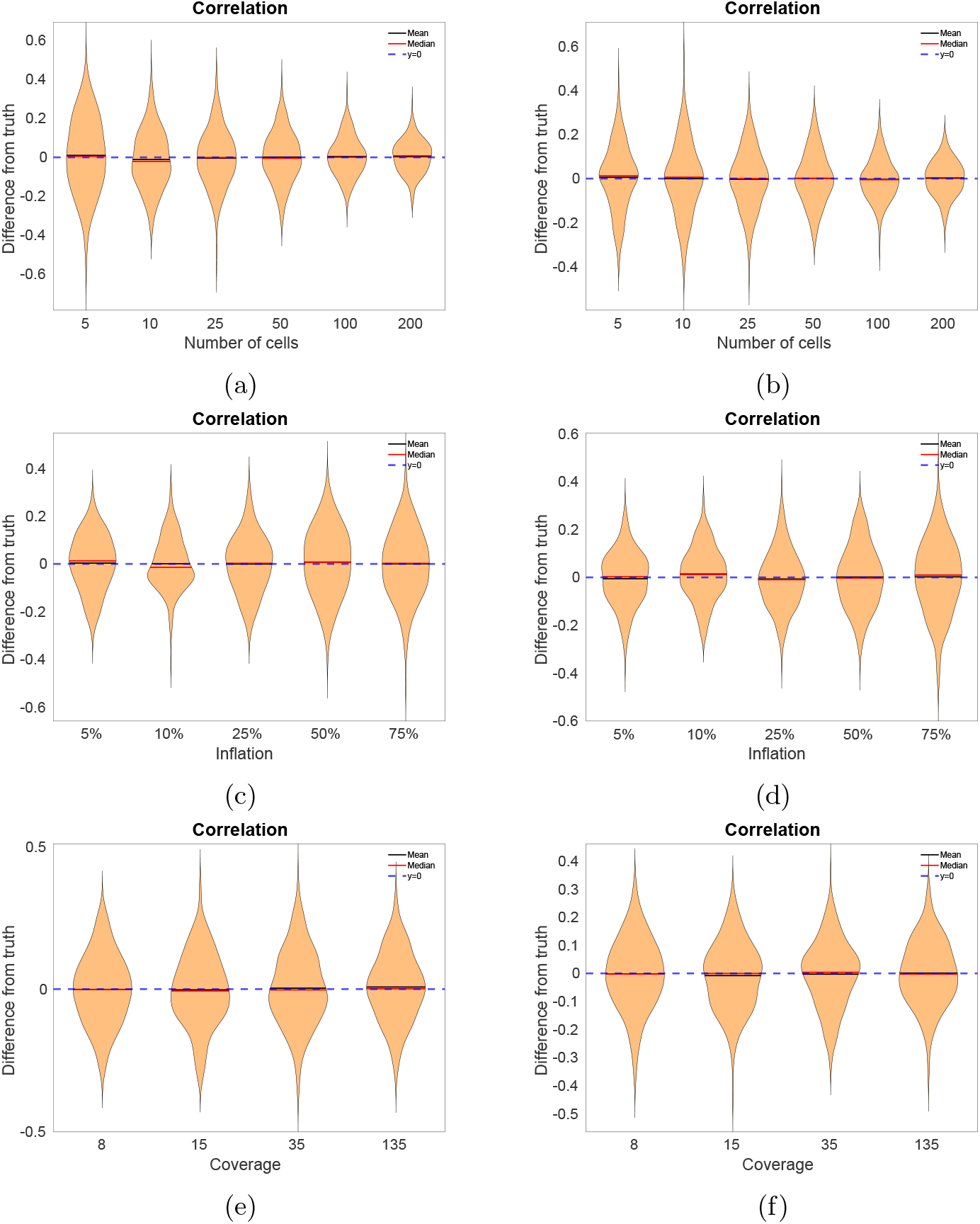
Estimation performance for SCRaPL based on synthetic data. The data generation mechanisms used for each experiment are described in *Methods*. Difference between true and the inferred correlation parameters *ρ*_*j*_ in synthetic data sampled from the model as a function of: (2a) the number of cells (experiment 1), (2c) the average gene inflation rate *π*_*j*_ (experiment 2), (2e) the average coverage (experiment 3). Difference between true and inferred correlation parameters *ρ*_*j*_ in synthetic data partly sampled from the model and partly from a deep generative model described in [18] as a function of: (2b) the number of cells (experiment 7), (2d) the average gene inflation rate *π*_*j*_ (experiment 8), (2f) the average coverage (experiment 9). Results for the remaining experiments are provided in Supplementary Figures S2-S6. In all cases, violin plots show the distribution across genes.

As a final test of more severe model mismatch, we evaluated predictive performance in a scenario where we retained the same noise model, but replaced the latent multivariate Gaussian distribution by expression rates inferred using a variational auto-encoder (scVI, [18]) that was trained on the scRNAseq data from [4] (*Methods*). Despite the model mismatch, we observed good estimating performance for *ρ*_*j*_ across a range of simulation parameters (Figures 2b,2d and 2f).

### 2.3 SCRaPL improves the power to identify associations between methylation and expression in mouse embryonic stem cells

We next considered a single cell multi-omics dataset generated by the scNMT-seq protocol [6]. Samples correspond to mouse embryonic stem cells (mESCs) at four developmental stages (embryonic days 4.5, 5.5, 6.5 and 7.5), which comprise the exit from pluripotency and primary germ layer [19] (see a brief description in the *Methods*). Here, all developmental stages were analysed together as a single group of samples. For each gene, matched methylation data was derived for protein coding promoters within 5kbp windows (chromatin accessibility data was also available, but excluded from our analysis). After quality control, the resultant dataset contained 9480 genes and average of 487 cells (*Methods*).

As a comparison to the correlation test implemented in SCRaPL, we considered the Pearson’s correlation test (*Methods*). The latter has been widely used when analysing single cell multi-omics data (e.g. [4, 6]), but does not take into account technical noise. Expression-methylation associations are retrieved as significant by controlling EFDR and FDR to 10%, respectively. We observe that SCRaPL retrieves approximately 3 times more associations compared to Pearson’s testing (198 versus 68; see Supplementary Table S1). This is despite the use of a more conservative correlation threshold *γ* = 0.205 in SCRaPL (which is calibrated via permutations, see *Methods*). To visually assess the results, we use volcano plots: Figure 3a shows a Bayesian volcano plot (median posterior correlation against the negative logarithm of the posterior probability in (9)), while Figure 3b is a standard volcano plot showing Pearson correlation versus negative logarithm of the associated *p*-value. While a substantial fraction of significant features are shared, there are many features on which both methods disagree, indicating a qualitative difference between the results provided by the two methods. We explore those cases in the next sections.

**Figure 3:**
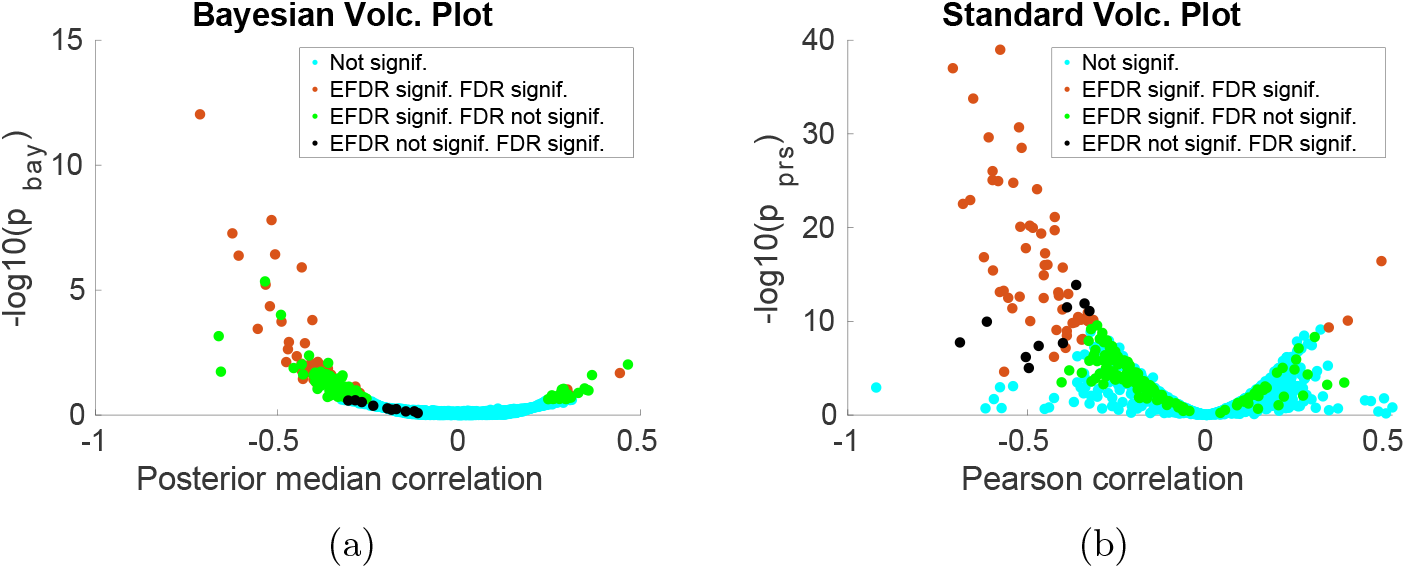
Bayesian 3a) and frequentist 3b versions of volcano plots for 5kbp window around TSS for [19] data. SCRaPL correlation and cut-off thresholds for choosing genes are chosen to be *γ* = 0.205 and *α* = 0.765 respective. Each dot represents a feature and is marked with different color depending the model that labels it important.

While the ability of SCRaPL to detect larger numbers of associations is certainly an encouraging feature, it is essential to characterize whether this is due to greater power, or simply to a greater vulnerability to false positives. Determining empirically the false positive rate is challenging as access to ground truth correlation values for each gene is impossible. To address these issues, we proceed pragmatically by constructing negative control data sets in which observations of methylation and expression values for a particular feature in different cells are randomly permuted. This will destroy any correlation structure between the two quantities, so that features detected as significant in negative control data can be considered as false positives. Here, we constructed 5 negative control datasets. For all negative controls, SCRaPL and Pearson’s testing only detected a handful of associations, consistently less than for the original data (Supplementary Table S1). These results suggest that both methods can detect genuine associations, albeit Pearson’s testing being less powerful.

### 2.4 SCRaPL associations are influenced by methylation coverage and are robust to outliers

As seen in Figures 3a and 3b, there is some discrepancy in terms of the associations detected by both Pearson correlation and SCRaPL. It is therefore natural to wonder to what extent the signals detected by the two methods are different. From the modelling perspective, there are two major differences: first, SCRaPL considers noise models which allow overdispersion and take into account coverage in the methylation data. This should make SCRaPL associations less vulnerable to outlier expression values or to methylation measurements with low coverage. Secondly, SCRaPL includes zero inflation in its expression model, and can therefore attribute to dropout some measurements of zero expression should the evidence dictate so. In the rest of this section, we present some empirical evidence that indeed observed these benefits in our real data analysis.

We consider the set of associations which are called as significant by at least one method, and split it into 3 categories: agreement between predictions, association labeling as significant by SCRaPL, but not Pearson, and vice-versa. We then analyze these three sets attempting to detect common patterns, discussing some examples to substantiate our findings.

Features for which Pearson and SCRaPL agree tend to have high number of observations, high coverage and small number of zeros in case of expression. An example feature called as significant by both methods is in Figure 6a.

To gain more insight on the factors driving SCRaPL inferences it is interesting to focus on associations, whose significance differs between the two methods. An example of an association detected by Pearson but not SCRaPL is in Figure 4b. As we can see, we have a large fraction of zero expression values with very low methylation coverage. As a result, SCRaPL, while placing most of the posterior mass over negative correlation values, cannot confidently exclude the possibility of no correlation. This example perfectly illustrates that divergences between SCRaPL and Pearson are often driven not by expected values, but by the fact that SCRaPL additionally performs uncertainty quantification on its results.

**Figure 4:**
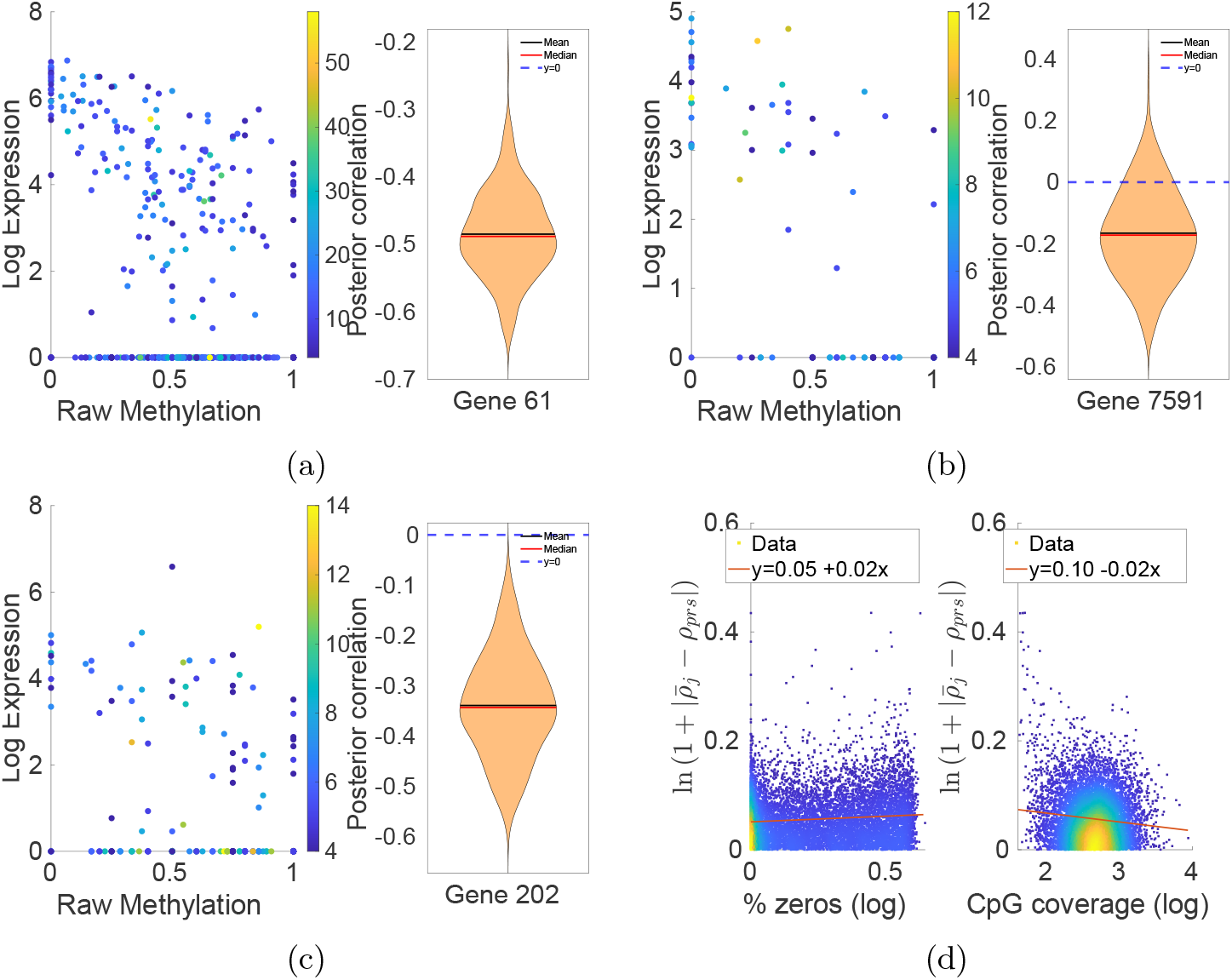
Where 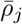 and *ρ*_*prs*_ are posterior mean and Pearson correlation for gene *j*. Examples that indicate SCRaPL’s behavior compared to Pearson correlation in micro and macro scale. In all figures apart from 4d the left part is a scatter plot of feature’s raw data colorcoded by CpG coverage with normalized expression in the *log*(1 + *x*) scale. On the right, we see the posterior for the same feature as inferred by SCRaPL. (6a)Agreement between SCRaPL and Pearson. (4c) Marked as important only by SCRaPL. (4b) Marked as important only by Pearson. In figure 4d we have scatter plots of distance between posterior mean and Pearson correlation as a function of inflation and coverage

An example of an association deemed significant by SCRaPL, but not by Pearson, is in Figure 4c. In this case, we tend to have medium to high expression, high number of observations and good coverage. However, Pearson correlation remains below detection levels due to observations with low expression. This is an example where SCRaPL can be particularly beneficial, since the noise model can better capture potential effect of zero inflation.

To provide a more quantitative, global explanation of the differences between SCRaPL and Pearson, we regress the absolute difference in inferred correlation against methylation coverage and percentage of zero counts for each gene across all cells. The resulting regressions, shown in Figure 4d, demonstrate a weak but consistent effect of both forms on noise, confirming that differences between the two methods are more prominent in noisier situations where methylation coverage is low or sparsity is high.

### 2.5 SCRaPL identifies biologically meaningful epigenetic regulation in early mouse gastrulation

SCRaPL identified a series of statistical associations between epigenomic and transcriptomic layers by addressing noise specifically encountered in single cell multiomics data. Markers with strong correlation were further investigated, using Gene Set Enrichment Analysis (GSEA) to establish links with biological phenomena observed in early embryogenesis or gene promoter methylation. DAVID [20] identified a mixture of developmental and house-keeping processes. Among processes tied to development, we see in utero embryonic development and angiogenesis. Whereas house-keeping ones include positive regulation of cell proliferation and regulation of transcription. For more information, the reader is directed in supplementary figure s13.

Apart from GSEA we looked at gene families like Dnmt and Dppa whose role has been studied [19, 21, 22] due to their known links to epigenetic regulation embryonic development respectively. Developmental pluripotency markers like Dppa2, Dppa4 and Dppa5a presented strong negative regulation between methylation and gene expression. From the Dna methyltransferases family only Dnmt3l had significant correlation. For further information please refer to supplementary figures S12-S14.

## 3 Conclusion

Single cell multi-omics sequencing technologies are rapidly becoming an important tool to understand epigenetic regulation for individual cells in complex biological processes, such as early embryo development. However, analysis of such data still presents a major bottleneck, due to the high-dimensionality, sparsity and heterogeneous noise affecting them. In this paper, we argued that the introduction of noise-aware approaches is fundamental in developing the field of single-cell multi-omics. We introduced SCRaPL, a Bayesian approach to perform perhaps the most basic and common multi-omics analysis, the discovery of correlative associations between different data modalities. By employing dedicated noise models in a latent-Gaussian framework, SCRaPL achieves more powerful and more robust results than simple analyses based on Pearson correlation, which is by far the most widespread tool currently used.

In our analyses, for each gene, matching methylation data was quantified using pre-defined promoter regions. This appears to be a reasonable demonstration of the tool; however it should be pointed out that SCRaPL could also be used to test associations between unannotated regions and expression of specific genes. Additionally, while our analyses have concerned associations between expression and DNA methylation, it is straightforward to extend the method to analyze correlations between different omic modalities, for example chromatin accessibility and gene expression [10], [23].

The Bayesian hierarchical framework employed by SCRaPL also offers a template for the application of more complex analysis techniques (such as clustering, dimensionality reduction and network inference) to multi-omics data. In all analyses, we expect that consistent handling of noise will improve robustness and biological significance. However, as with most Bayesian methods, SCRaPL does suffer from a higher computational burden, particularly when compared with extremely simple analyses, such as Pearson correlation. Extension of noise-aware Bayesian methods to different single-cell multi-omics analyses will require the adoption and evaluation of more efficient computational inference techniques, such as variational inference [24].

## 4 Methods

### 4.1 A Bayesian hierarchical framework for noisy single cell multi-omics data

SCRaPL implements a Bayesian hierarchical approach that is tailored to the data generated by single cell multi-omic assays. Here, we assume that matched data is available for two molecular phenotypes, but our formulation is flexible and can be expanded to include additional layers. A graphical representation for the model implemented in SCRaPL is provided in Figure 5. As described in Figure 1, the distribution of a latent vector **X**_*ij*_ is used to capture the association across molecular layers. For each cell *i* (∈ {1, … *I*}) and gene *j* (∈ {1, … *J*}), the latter is given by

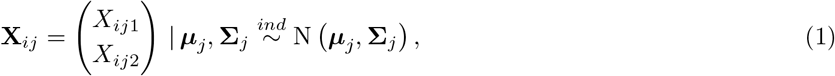

where

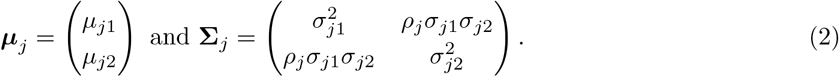

**Figure 5:**
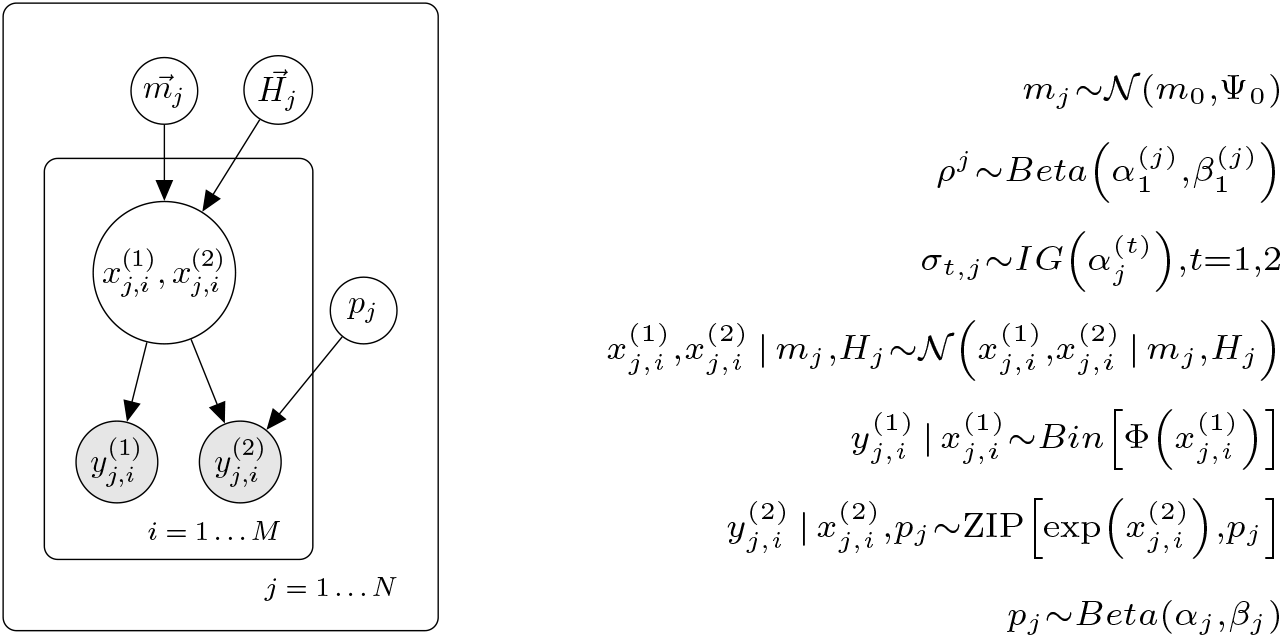
Graphical model implemented in SCRaPL

**Figure 6:**
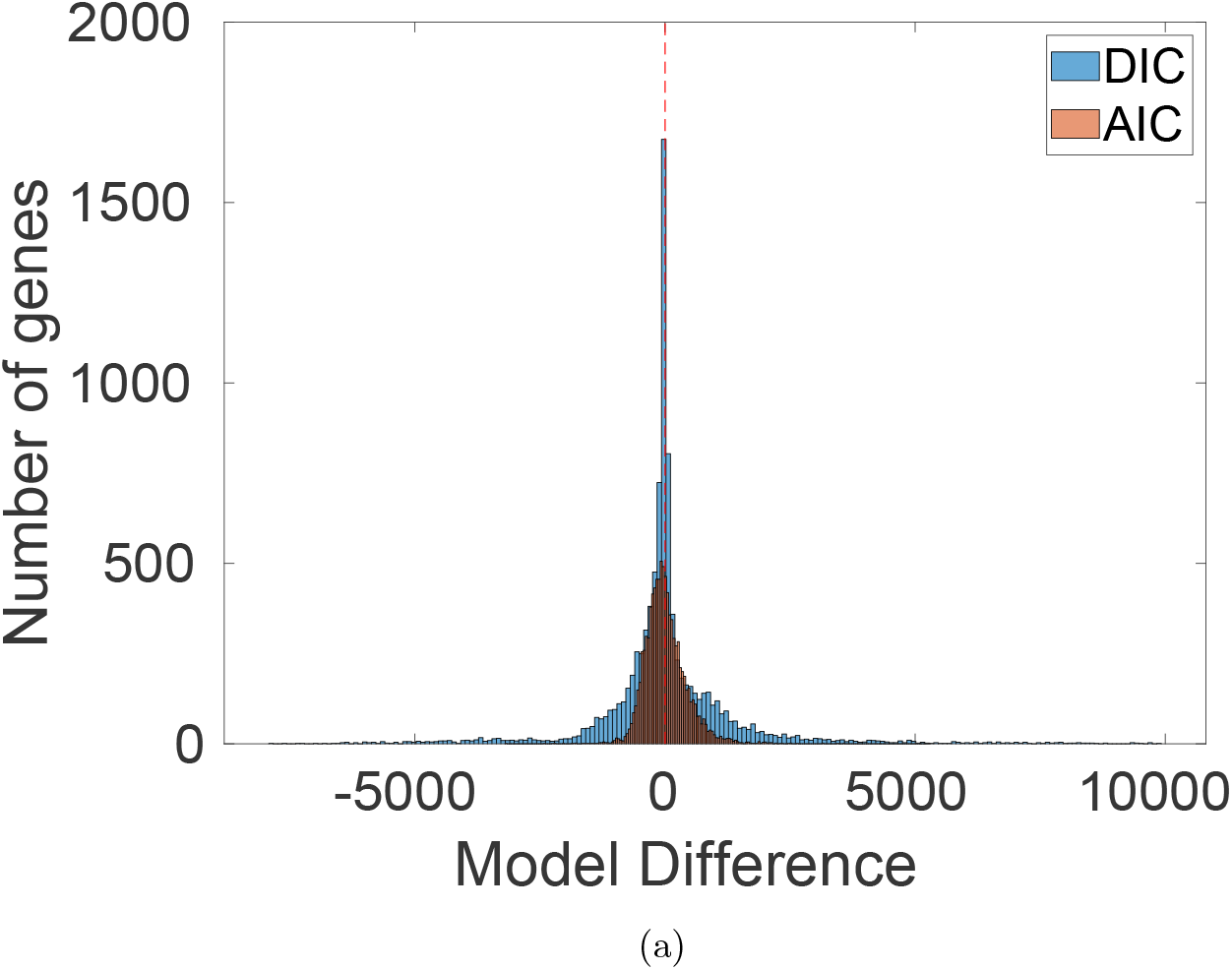
Difference between model with and without inflation. The more negative the difference, the stronger the evidence in favor of the model with zero inflation is and vice versa. For comparison AIC is also plotted in red.

In this formulation, we assume independence across all genes. Moreover, different noise models are then assigned to each molecular layer based on the properties of the associated data. In particular, the noise models used by SCRaPL in the context of RNA expression and DNAm data are described below.

#### RNA expression noise model

Let *Y*_*ij*1_ be a random variable representing the number of read-counts observed for each cell *i* and gene *j*. Conditional on the value of the latent variable *X*_*ij*1_, we use an exponential link function and assume that

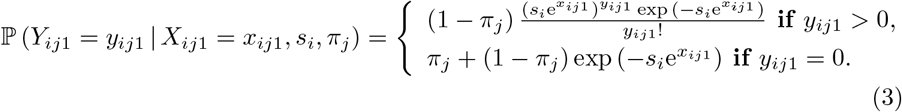

The latter corresponds to a zero-inflated Poisson (ZIP) model, where *s*_*i*_ (*>* 0) is a cell-specific scaling factor that accounts for global differences across cells (e.g. due to sequencing depth) and *π*_*j*_ (∈ [0, 1]) represents a zero-inflation parameter. In practice, we infer scaling factors *s*_*i*_ using scran [15] and use them as known offsets. If *π*_*j*_ = 0, (3) reduces to a Poisson model. The need for a zero-inflation component is a matter of debate for scRNA-seq data [13] and may depend on the experimental protocol used to generate the data. For each gene, here we quantify the evidence in favour of zero-inflation using.

#### DNAm noise model

For each cell *i* and gene *j*, let *n*_*ij*_ be the number of CpG sites for which DNAm reads were obtained. These capture differences in coverage across cells and genes. The conditional model for the number of methylated CpG sites *Y*_*ij*2_ is then assumed to follow a binomial distribution such that

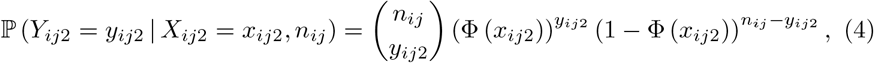

where Φ(*·*) denotes a probit link function.

#### Parameter interpretation

To aid the interpretation of each model parameter, mean and variance expressions are derived for the noise models introduced above after integrating out the distribution of the latent vector **X**_*ij*_ (see Supplementary Section S8). In both cases, *µ*_*j*1_ and *µ*_*j*2_ control the overall RNA expression and DNAm values for the population of cells under study. Moreover, *σ*_*j*1_ and *σ*_*j*2_ capture the excess of variability (overdispersion) that is observed with respect to the baseline noise model. Finally, *ρ*_*j*_ captures the latent correlation between molecular layers.

### 4.2 Prior specification

A popular prior choice for covariance matrices is the inverse Wishart distribution. However, this has been shown to bias correlation coefficients depending whether marginal variances are small or large [25]. In [26] they use a separation strategy to decouple correlation from marginal variances. Then, one approach is to keep Inverse gamma for marginal variances and use uniform priors. Our prior specification for **Σ**_*j*_ is based on the parametrization introduced in (2), with independent priors assigned to all gene-specific parameters. Our prior specification is given by

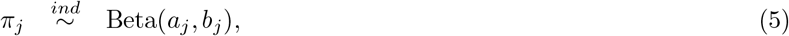

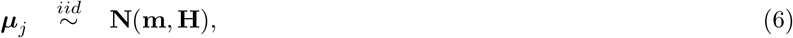

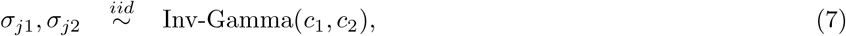

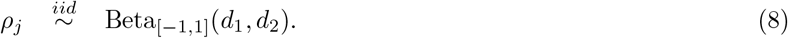

In (8), the prior for *ρ*_*j*_ corresponds to a four-parameter Beta distribution, whose support has been scaled to be [−1, 1].

### 4.3 Implementation

As the posterior distribution associated to the model above does not have a closed analytical form, inference is implemented using a mixture of Hamiltonian Monte Carlo (HMC) [27] and Gibbs Sampler [28]. Apart from latent mean parameters ***µ***_*j*_ which is sampled with Gbbs, the remaining parameters are sampled using HMC. HMC updates were implemented and tunned using the tuneSampler Matlab library [29].

For all the analyses shown in this article, we obtained 1500 samples from this algorithm and discarded the first 1050 iterations (burn-in) before estimating model parameters. Convergence is monitored using the Gelman-Rubin criterion [30].

### 4.4 A probabilistic rule to detect statistically significant associations across layers

SCRaPL identifies genes with statistically significant correlation across molecular layers (e.g. RNA expression and promoter DNAm) based on the posterior distribution of gene-specific latent correlation parameters *ρ*_*j*_. Our decision rule depends on whether the posterior mass for |*ρ*_*j*_| is concentrated around high values. As in [16], this is quantified by the following tail posterior probabilities

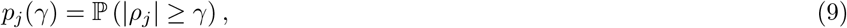

where *γ* (*>* 0) denotes a minimum correlation threshold. If *p*_*j*_(*γ*) is greater than a probability threshold *α*, a statistically significant correlation is reported for gene *j*.

Suitable values for *γ* and *α* could be chosen using different approaches. In principle, *γ* can be fixed a priori by the user. Instead, we adopt a data-driven approach based on the distribution of gene-specific posterior estimates obtained for |*ρ*_*j*_| using negative control datasets (Supplementary Section S4). Such distribution can be used to quantify the strength of correlation estimates that can be expected by chance for a given sample size and sequencing depth. As a default choice, we select *γ* to match the 90% quantile of the distribution described above. For a fixed value of *γ*, a grid search is used to select *α* according to a target Expected False Discovery Rate (EFDR) [17]. The latter is defined as

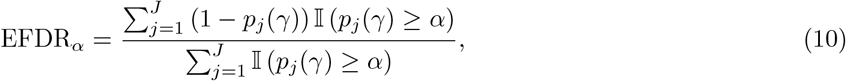

where 𝕀 (*A*) = 1 if *A* is true, 0 otherwise. Our default target EFDR is equal to 10%.

### 4.5 Current approach based on Pearson’s correlation

To date, single cell multi-omics analyses have primarily used the Pearson’s correlation coefficient *r* to quantify associations between different types of molecular data (e.g. [4][6]). These estimates are directly derived from the observed data and do not assume a specific noise model. As the input for this calculation, gene expression data is typically normalised (e.g. using scran [15]) and subsequently log-transformed after adding a pseudocount.

Based on these estimates, statistically significant correlations are selected by contrasting the hypotheses *H*_0_ : *r* ≤ *u* and *H*_1_ : *r* ≥ *u*, for me threshold *u*. To control the False Discovery Rate (FDR) across genes, the Benjamini-Hochberg correction [31] is typically used.

### 4.6 Comparing between alternative models

SCRaPL is a noise-aware approach designed to deal with different types of multiomics data. Dealing with data is achieved by incorporating likelihoods tailored to data produced by current and future sequencing technologies. With the ongoing debate on the importance of zero-inflation [13] still open, a natural question that arises is how to choose models that take into account zero-inflation with others that do not. In particular, how to decide if Poisson or zero-inflated Poisson is more appropriate for a specific gene. Since posterior model samples are available, we do model selection using Deviance Information Criterion (DIC) [14]. DIC is a method for assessing goodness of fit while penalizing large effective numbers of parameters between alternative models and it is estimated using 11.

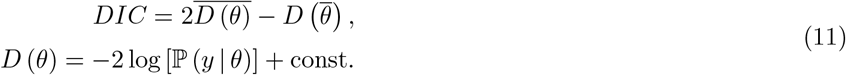

Where 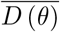 and 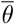 is the expectation of *D*(*θ*) and *θ* respectively wrt. *θ*. Essentially DIC assign a score to each alternative with lower DIC values indicating more preferable models. To justify the use of zero-inflation in SCRaPL we fit the zero-inflated and the standard Poisson in the methylation/expression 2500kbp promoters of [6]. For around 60% of the genes in the dataset, DIC favors zero inflation.

### 4.7 Data aggregation

To aggregate methylation/accessibility data from different cells we follow a window based approach. Reads are mapped using the GRCm38 mouse genome (accession number GSE56879). For more information, the reader is directed to the appendix. As we would also like to understand how our window choices affect our results, especially when looking at enhancer regions where information tends to be more sparse, we consider multiple windows.

When looking at promoter regions in methylation/expression data in both datasets, a window of ±2.5kbp worked well.

### 4.8 Generating synthetic data

The first set of data were generated using SCRaPL’s generative model. We designed three types of experiments to asses estimation performance as a function of the number of cells, ZI for the expression data and methylation coverage. These were combined with different types of underlying correlation structure:

- **Experiment 1**: varying numbers of cells (*n* ∈ *{*5, 10, 25, 50, 100, 200*}*) and correlation values sampled from a Beta distribution (*ρ*_*j*_ ∼ Beta(15, 15)).
- **Experiment 2**: varying ZI rate (*ρ*_*j*_ ∈ *{*0.05, 0.10, 0.25, 0.50, 0.75*}*) and correlation values sampled from a Beta distribution (*ρ*_*j*_ ∼ Beta(15, 15)).
- **Experiment 3**: varying methylation coverage (*n*_*ij*_ sampled from Uniform distributions with ranges given by [5, 10], [10, 20], [20, 50] and [50, 250]) and correlation values sampled from a Beta distribution (*ρ*_*j*_ ∼ Beta(15, 15)).
- **Experiment 4**: varying numbers of cells (*n* ∈ *{*5, 10, 25, 50, 100, 200*}*) and constant correlation across genes (*ρ*_*j*_ = 0.7).
- **Experiment 5**: varying ZI rate (*ρ*_*j*_ ∈ *{*0.05, 0.10, 0.25, 0.50, 0.75*}*) and constant correlation across genes (*ρ*_*j*_ = 0.7).
- **Experiment 6**: varying methylation coverage (*n*_*ij*_ sampled from Uniform distributions with ranges given by [5, 10], [10, 20], [20, 50] and [50, 250]) and constant correlation across genes (*ρ*_*j*_ = 0.7).

In all cases, latent means and standard deviations were set as *µ*_*j*1_ = 1, *µ*_*j*2_ = 4, *σ*_*j*1_ = 2 and *σ*_*j*2_ = 3. Unless otherwise stated, our simulations were based on: *M* = 60 cells, *N* = 300 genes, 10% ZI rate for the expression data (*π*_*j*_ = 0.10) and an average methylation coverage (*n*_*ij*_) equal to 275 across cells and genes.

A second set of experiments was designed to evaluate the performance of SCRaPL under model mismatch. For this purpose, instead of generating latent expression values *X*_*ij*1_ from a normal distribution, these were generated using scVI [18] (Supplementary Figures S2-S6). Under this setup, the following experiments were performed:

- **Experiment 7**: varying numbers of cells (*n* ∈ *{*5, 10, 25, 50, 100, 200*}*) and correlation values sampled from a Beta distribution (*ρ*_*j*_ ∼ Beta(15, 15)).
- **Experiment 8**: varying ZI rate (*ρ*_*j*_ ∈ *{*0.05, 0.10, 0.25, 0.50, 0.75*}*) and correlation values sampled from a Beta distribution (*ρ*_*j*_ ∼ Beta(15, 15)).
- **Experiment 9**: varying methylation coverage (*n*_*ij*_ sampled from Uniform distributions with ranges given by [5, 10], [10, 20], [20, 50] and [50, 250]) and correlation values sampled from a Beta distribution (*ρ*_*j*_ ∼ Beta(15, 15)).
- **Experiment 10**: varying numbers of cells (*n* ∈ *{*5, 10, 25, 50, 100, 200*}*) and correlation values sampled from a Uniform distribution (*ρ*_*j*_ ∼ U(−0.8, −0.6)).
- **Experiment 11**: varying ZI rate (*ρ*_*j*_ ∈ *{*0.05, 0.10, 0.25, 0.50, 0.75*}*) and correlation values sampled from a Uniform distribution (*ρ*_*j*_ ∼ U(−0.8, −0.6)).
- **Experiment 12**: varying methylation coverage (*n*_*ij*_ sampled from Uniform distributions with ranges given by [5, 10], [10, 20], [20, 50] and [50, 250]) and correlation values sampled from a Uniform distribution (*ρ*_*j*_ ∼ U(−0.8, −0.6)).

As before, unless otherwise stated, our simulations were based on: *M* = 60 cells, *N* = 300 genes, 10% ZI rate for the expression data (*π*_*j*_ = 0.10) and an average methylation coverage (*n*_*ij*_) equal to 275 across cells and genes.

## Acknowledgments

Our work was supported in part by the EPSRC Centre for Doctoral Training in Data Science, funded by the UK Engineering and Physical Sciences Research Council (grant EP/L016427/1 to Christos Maniatis) and the University of Edinburgh. We would also like to thank Chantriolnt-Andreas Kapourani, Michalis Michaelides for their valuable comments and discussion.

