## Supplementary Material for "SCRaPL: hierarchical Bayesian modelling of associations in single cell multi-omics data"

### Supplementary Materials

Christos Maniatis

February 2020

#### S1 Data preprocessing

Data preprocessing is a vital step prior to inference and is split into two parts, quality control(QC) and integration. Quality control ensures that individual components and joint data satisfy a minimum set of requirements before inference. As single cell sequencing data are stored into different files and formats, appropriate steps need to be considered to bring them in a format understandable by the model. These steps are part of the integration and involve aggregating information for each region in each molecular layer and later joining information between different epigenetic marks using a unique label linked to the region. In this section we give a detailed presentation of integration and quality control steps justifying some of our choices. In many cases integration steps take place between quality controls, so it is natural to present QC and integration steps in the order they appear.

For certain genomic regions when methylation/accessibility data are collected using sc-NOME-seq (Pott, 2017) it is common to have multiple readings. The first step is to average these readings and binarise them in order to get an index for each cytosine. Then using region locations and cell label, methylation and accessibility readings are accumulated into one file. Essentially the script iterates through each cell and aggregates methylated cytosines and total number of cytosines using a window based approach for each region. In this step, user feeds the script with the window length and the minimum coverage a region needs to have in order not to be discarded. Observations in regions with low coverage are marked with $-1$ and are discarded later in the pipeline. These steps take raw binarised data and yield methylation/accessibility or both for genomic region of interest and cell label.

Single cell RNA expression is initially filtered based on the total number of counts per cell, minimum number of expressed genes per cell and distribution of counts. Using scran we then estimate a unique normalization constant for each cell. In this step we take raw expression data, filter low quality cells and estimate per cell normalization constants.

At this point we have aggregated methylation/accessibility data and expression counts with the corresponding normalization constant. For each observation we have a pair of labels indicating genomic region and cell. These labels are used to integrate molecular layers. Then regions with low methylation/accessibility coverage marked with $-1$ are discarded. The final step requires to pass genomic regions through a joint QC control. There regions with less than 5 reads, zero variance in each component and percentage of expression zeros above 80% are removed. The threshold for observations was set to 5 because for less reads the the model heavily relied on prior. The 80% threshold for expression zeros was set to balance between available non-zero readings in each feature and keeping as many genomic regions as possible. Since correlation is highly dependent on components variability, it vital to ensure positive variance and avoid inference problems.

The final step of preprocessing is to store the filtered epigenetic marks in a matrix. Independently of which pair of epigenetic marks we are dealing with, the convention we use is to place methylation pairs in the first two columns of the matrix and to store expression then. Hence if we have a methylation/accessibility pair, methylation will be first and accessibility second. Similarly in the case of accessibility/expression pair, accessibility will occupy the first 2 columns and expression will follow. Then we store numerical indices corresponding to unique genomic and cell labels, in order to guide the model to consider the correct observations and allow different number of observations per feature. Furthermore, three tables accompany the matrix The first one has normalization constants and cell labels, the second and third have actual cell and feature names. Scripts with all preprocessing and integration steps can be found here [1]. We proceed with the description of negative control pipeline.

[1] github.com/chrmaniatis/SCRaPL

### S2 Creating negative control datastes

Negative control datasets were created to understand model's robustness against false positives. The main idea is that if the model mainly detects false positives, then by shattering any correlation structure we expect to detect similar number of features. In this subsection we give a thorough description of the pipeline creating these datasets.

When constructing negative control data the non-missing methylation/accessibility along with expression values are permuted. To keep the analysis simple, we assume that the starting dataset has single cell methylation and expression data. In our analysis we assume that each component is stored in a separate tensor as depicted in S1a. Rows and columns of each tensor represent features and cells respectively. Missing methylation are denoted with black boxes. Scrambling cell labels in this setting essentially means that we column-wise permute tensors (without permuting its labels). As there are missing methylation values it is not possible to column-wise permute the entire tensor, because expression values originally paired with a missing methylation will end-up with non-missing ones changing dataset properties and making comparison difficult.

To bypass this issue we iterate through existing features, generating distinct permutation for each molecular layer. For epigentic marks with missing values (ie. methylation and accessibility) we scramble non-missing cells. With expression things are straight-forward as all values are in place but care needs to shown with normalization constants which are tied to specific cells. This is presented in figure S1b. Pseudocode for creating negative control data is presented in 1.

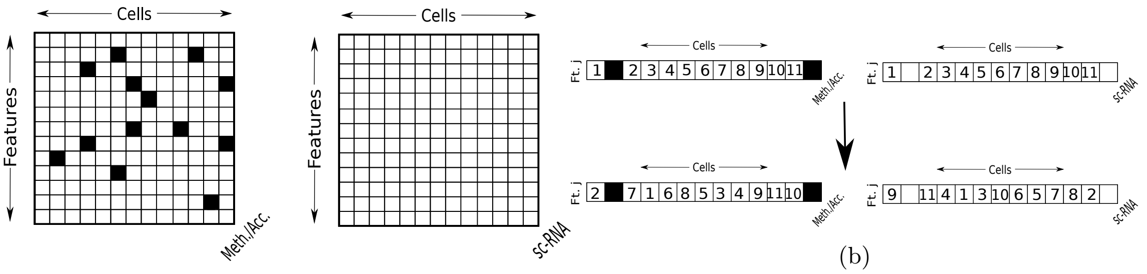


(a) (b)

Figure S1: (S1a) Sketch of tensors storing methylation/accessibility(left) and expression(right) respectively. Black boxes represent missing methylation/accessibility data. (S1b) Sketch of cell scrambling for a random feature j. It is important to note that despite cells are permuted at the end of scambling, cell labels remain fixed.

**Algorithm 1** Negative control dataset pipeline

**Input:** Dataset of interest.
**Output:** Negative control dataset

Load dataset of interest.
Compute the number of genomic features $N_g$.
**for** $ii = 1, 2, \ldots, N_g$ **do**
    Compute number of cells $N_c$ for that genomic region of interest.
    Define 2 vectors with all integers ranging from 1 to $N_c$ with dimension $1 \times N_c$.
    Compute two distinct random permutations for these vectors.
    Apply permutation the permutation in each epigentic mark.
Save updated data into a new file.

### S3 Synthetic Data

In experiments involving synthetic data, three sets of experiments were preformed. For the first set of experiment, samples from the prior were used. The second one involved samples unlikely to appear given the probabilistic graphical model describing the problem. Under prior correlation $\rho_j \sim Beta(15, 15)$, sampling a correlation value 0.7 would have probability below $\frac{1}{2000}$. In the third one we consider the case of model mismatch. In other words, we assume that our modeling assumptions are violated in an attempt to understand how robust is SCRaPL in case of a model mismatch. To achieve that, we use a variational auto-encoder to estimate a more realistic distribution for expression whose samples in turn are used to condition the generative model, to sample methylation. In building such data we have the freedom of choosing how to generate correlation. Hence we have two scenarios; one where correlation is simulated from a $Beta(15, 15)$ and another one where it is sampled from to $U[-0.8, -0.6]$ following the same logic to the first set of experiments. For each set of experiments we tested SCRaPL's inference accuracy under scenarios where model's hyper-parameters like cell numbers, methylation coverage and inflation were varied. For each gene and each experiment we tracked a plethora of parameters like inflation, latent means and covariance. For space economy reasons in the benchmarks section of the main text, only plots of latent correlations for set of experiments with unlikely generating parameters are included. For the sake of completeness, here we include the rest of the experiments that were omitted from the main text.

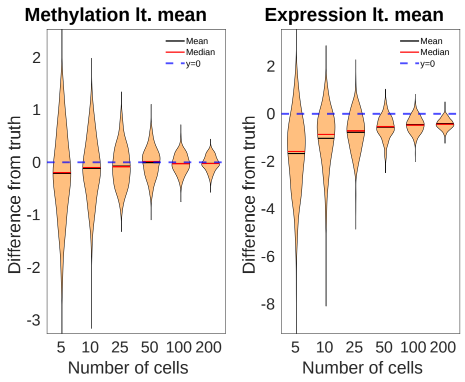


(a)

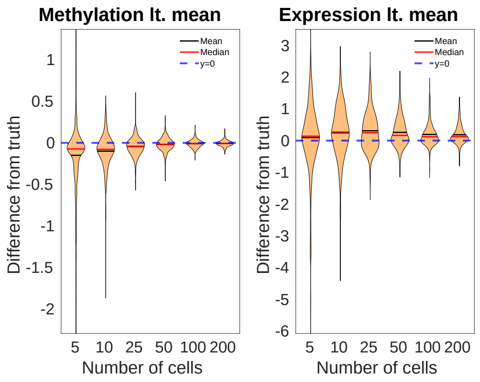


(b)

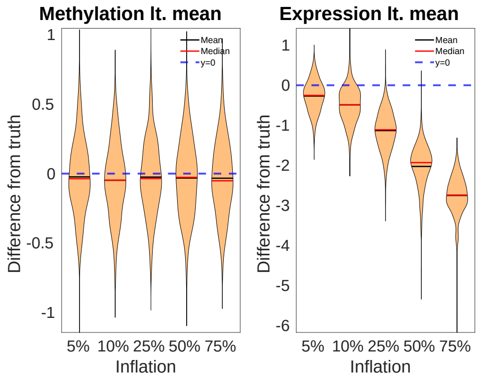


(c)

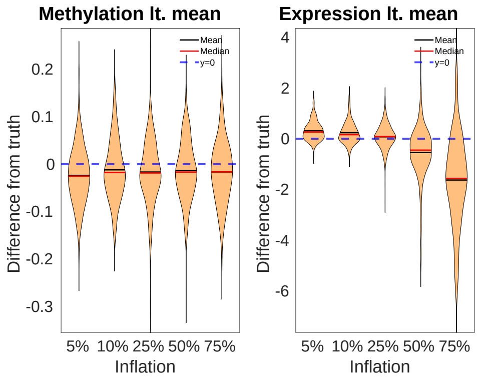


(d)

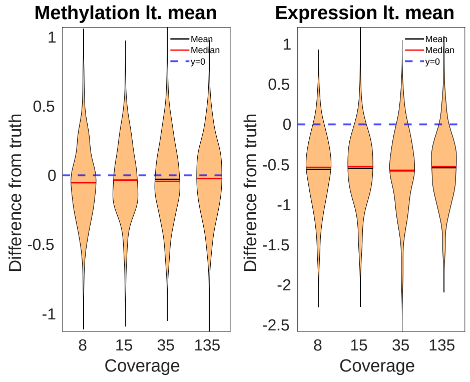


(e)

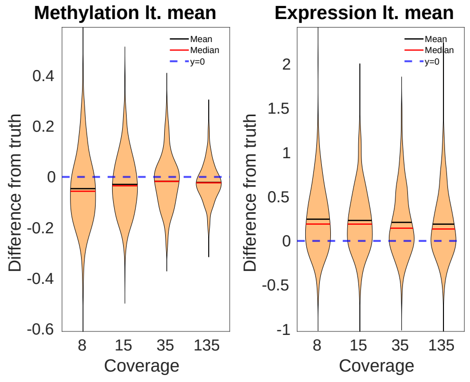


(f)

Figure S2: Difference between true and inferred latent means $\mu_j$ in synthetic data sampled from the model as a function of: (S2a) the number of cells (experiment 1), (S2c) average gene inflation rate $\pi_j$ (experiment 2), (S2e) the average gene coverage (experiment 3). Difference between true and inferred latent means $\mu_j$ in synthetic data partly sampled from the model and partly from a deep generative model described in Lopez et al. (2018) as a function of: (S2b)the number of cells (experiment 7),(S2d) gene inflation (experiment 8), (S2f)average gene coverage (experiment 9). In all cases, violin plots show the distribution across genes.

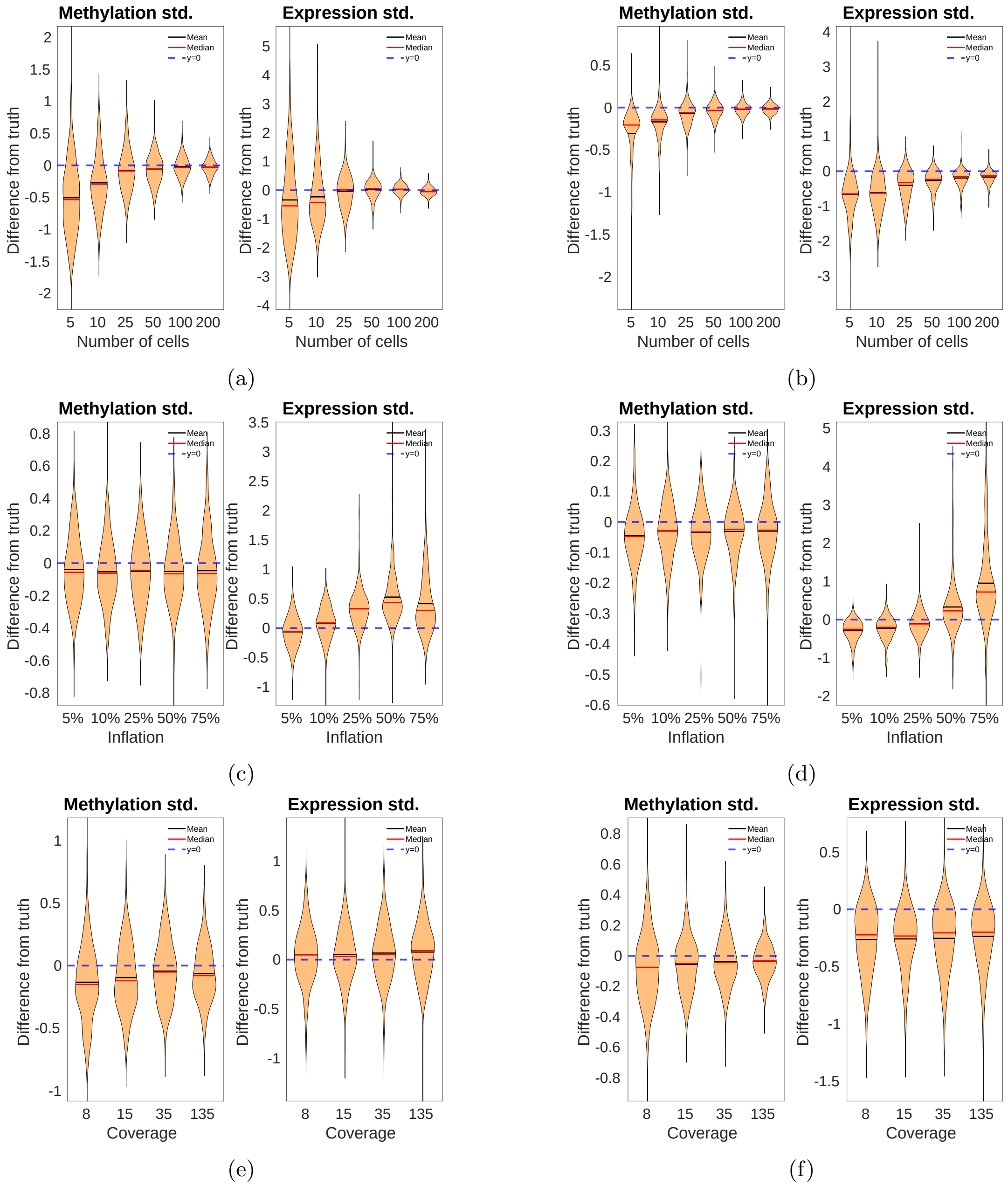


Figure S3: Difference between true and inferred latent standard deviations $\sigma_{j1}, \sigma_{j2}$, in synthetic data sampled from the model as a function of: (S3a) the number of cells (experiment 1), (S3c) average gene inflation rate $\pi_j$ (experiment 2), (S3e) the average gene coverage (experiment 3). Difference between true and inferred latent standard deviations $\sigma_{j1}, \sigma_{j2}$ in synthetic data partly sampled from the model and partly from a deep generative model described in Lopez et al. (2018) as a function of: (S3b)the number of cells (experiment 7),(S3d) gene inflation (experiment 8), (S3f)average gene coverage (experiment 9). In all cases, violin plots show the distribution across genes.

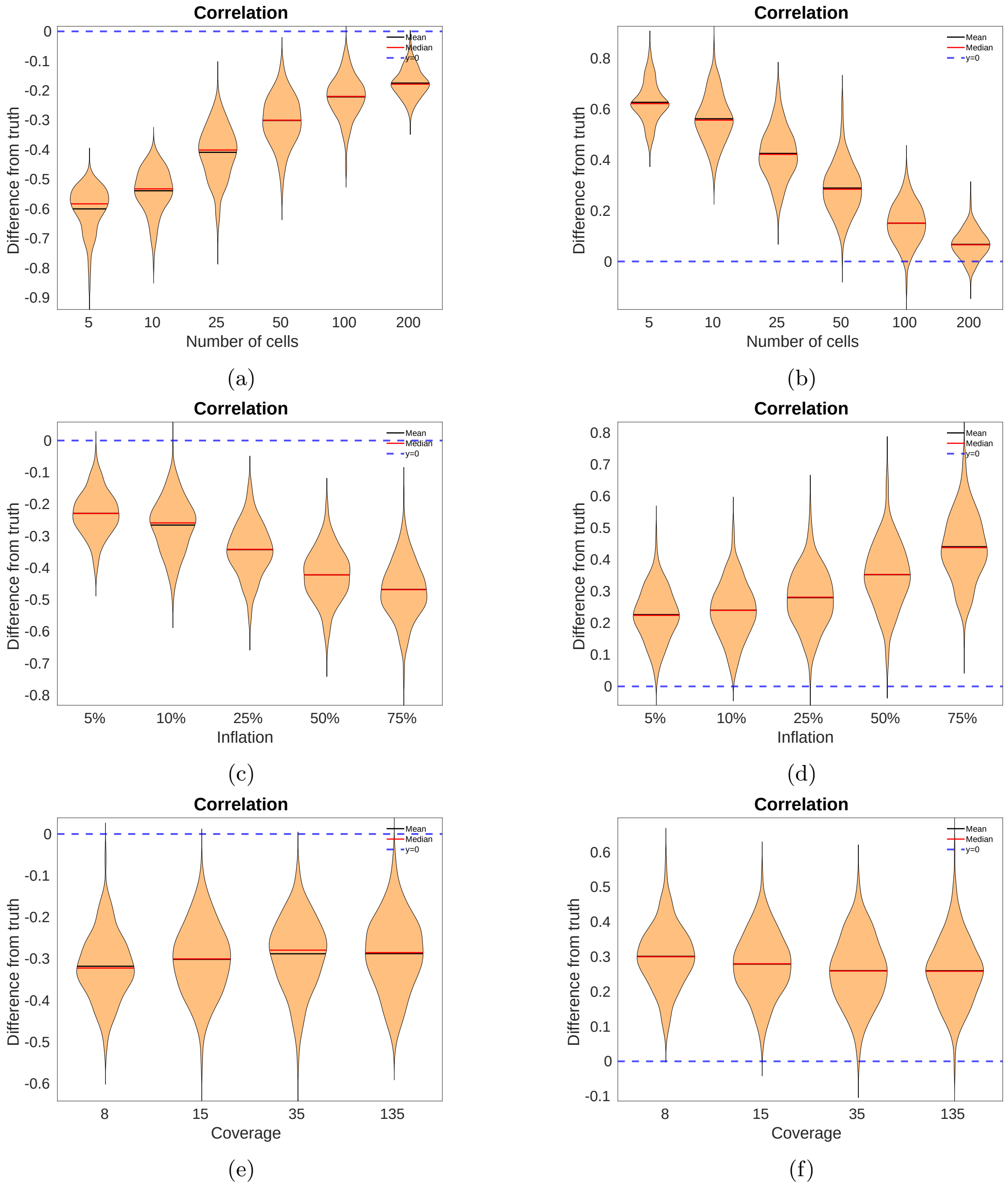


Figure S4: Difference between true and inferred latent correlations $\rho_j$ in synthetic data generated from the generative with correlation $\rho_j = 0.7$ model as a function of: (S4a) the number of cells (experiment 4), (S4c) average gene inflation rate $\pi_j$ (experiment 5), (S4e) the average gene coverage (experiment 6). Difference between true and inferred latent correlations $\rho_j$ in synthetic data partly sampled from the model with correlation $\rho_j \sim U(-0.8, -0.6)$ and partly from a deep generative model described in Lopez et al. (2018) as a function of: (S4b)the number of cells (experiment 10),(S4d) gene inflation (experiment 11),(S4f) average gene coverage (experiment 12). In all cases, violin plots show the distribution across genes.

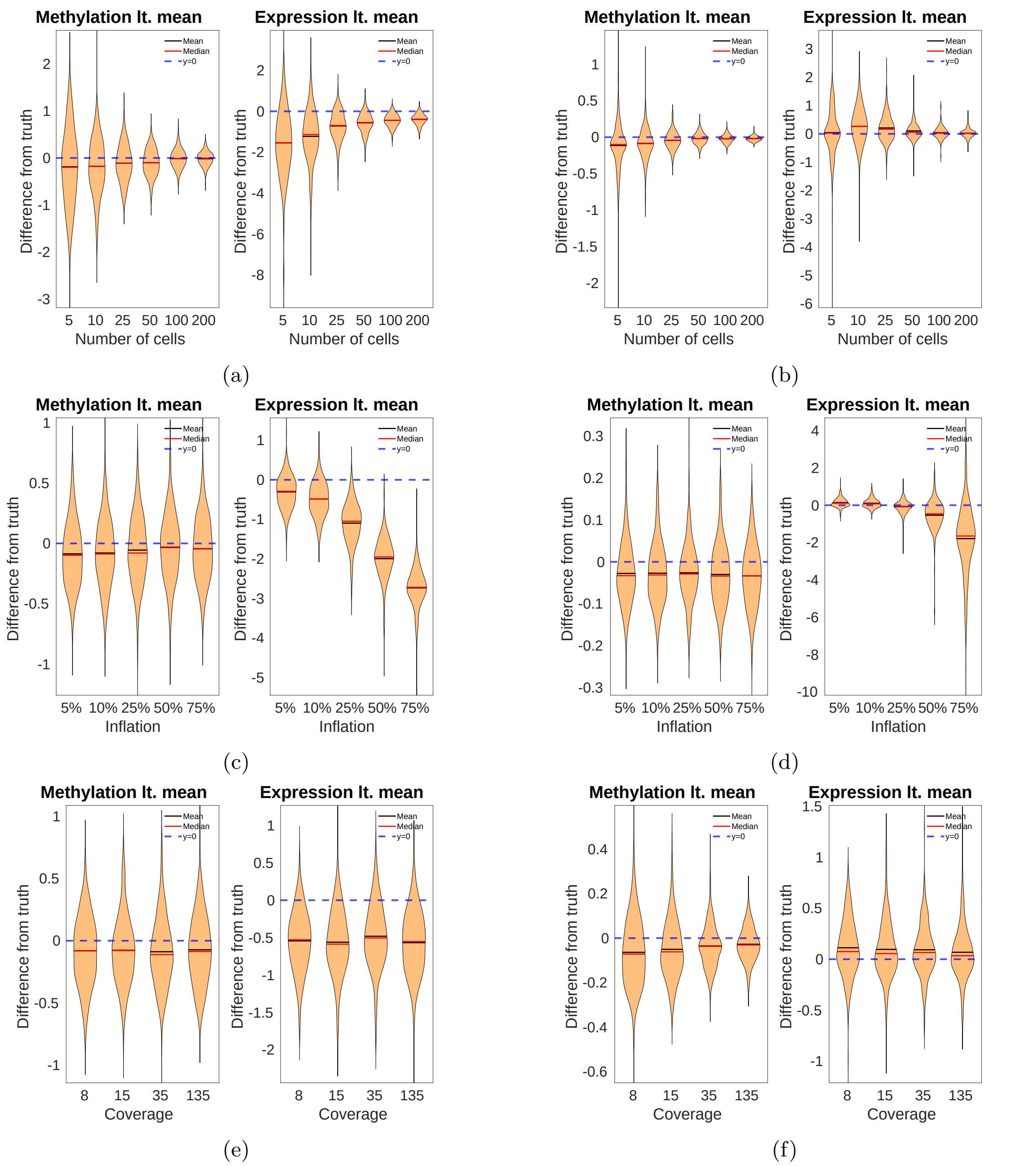


Figure S5: Difference between true and inferred latent means $\mu_j$ in synthetic data generated from the model with correlation $\rho_j = 0.7$ as a function of: (S5a) the number of cells (experiment 4), (S5c) average gene inflation rate $\pi_j$ (experiment 5), (S5e) the average gene coverage (experiment 6). Difference between true and inferred latent means $\mu_j$ in synthetic data partly sampled from the model with correlation $\rho_j \sim U(-0.8, -0.6)$ and partly from a deep generative model described in Lopez et al. (2018) as a function of: (S5b)the number of cells (experiment 10),(S5d) gene inflation (experiment 11),(S5f) average gene coverage (experiment 12). In all cases, violin plots show the distribution across genes.

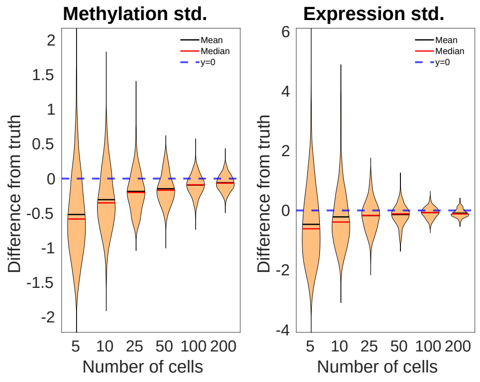


(a)

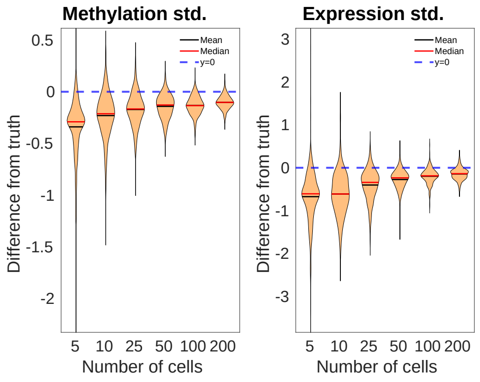


(b)

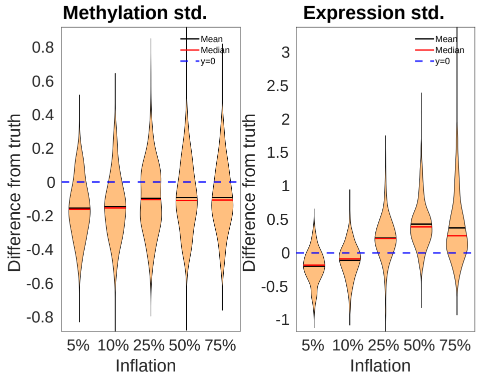


(c)

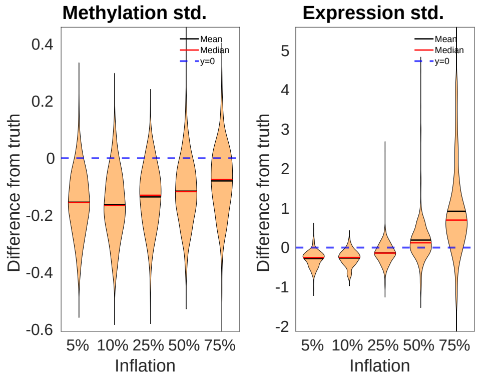


(d)

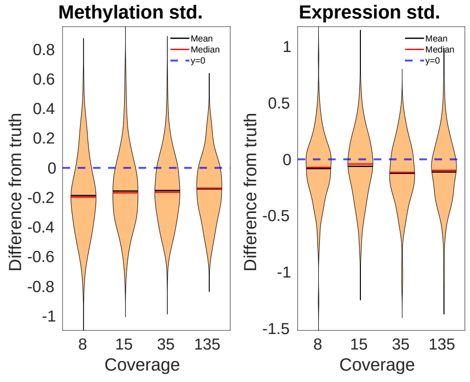


(e)

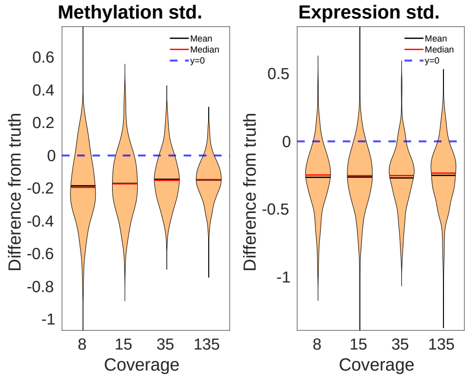


(f)

Figure S6: Difference between true and inferred latent standard deviations $\sigma_{j1}, \sigma_{j2}$ in synthetic data generated from the model with correlation $\rho_j = 0.7$ as a function of: (S6a) the number of cells (experiment 4), (S6c) average gene inflation rate $\pi_j$ (experiment 5), (S6e) the average gene coverage (experiment 6). Difference between true and inferred latent standard deviations $\sigma_{j1}, \sigma_{j2}$ in synthetic data partly sampled from the model with correlation $\rho_j \sim U(-0.8, -0.6)$ and partly from a deep generative model described in Lopez et al. (2018) as a function of: (S6b)the number of cells (experiment 10),(S6d) gene inflation (experiment 11),(S6f) average gene coverage (experiment 12). In all cases, violin plots show the distribution across genes.

It is evident from figures presented in this section that parameters with the highest impact are number of cells per feature and inflation, especially in the case of large generating correlations.

### S4 Negative control experiments

In this section, we demonstrate that SCRaPL detects significantly more features compared Pearson correlation while keeping false positives below accepted tolerance. To get these results threshold $\gamma$ was set to be 0.205 .

| | Orig. | Neg.1 | Neg.2 | Neg.3 | Neg.4 | Neg.5 |
|---|---|---|---|---|---|---|
| SCRaPL | 198 | 0 | 0 | 3 | 0 | 0 |
| Pearson | 68 | 0 | 0 | 0 | 0 | 0 |

Table S1: Number of detected features for original and negative control data with SCRaPL and Pearson correlation when $\gamma = 0.205$

From table S1, SCRaPL detects more features compared to Pearson correlation, while keeping false positives within accepted tolerance

#### S5 Choosing between priors

While building the model we had to find a systematic way of choosing between beta prior parameters. Since we did not want to favor neither positive nor negative correlations, we centered the prior at 0. Then to choose between a strict and a more liberal prior, we used original and negative control datasets. Posterior median correlation as a function of percentage of zeros encountered in each gene for strong and weak priors were compared with Pearson correlation against percentage of zeros. As we can see from Figure S7 the weak prior tends to be more liberal as the percentage of zeros increases. This behavior is also encountered in the plots when negative control data are used (Figure S8), potentially increasing the number of false positives. The strong prior on the other hand both in the true and negative control data yields more conservative results. Despite the fact that it could suppress true positive hits, it is preferred since the probability of finding a false positives is significantly below tolerance.

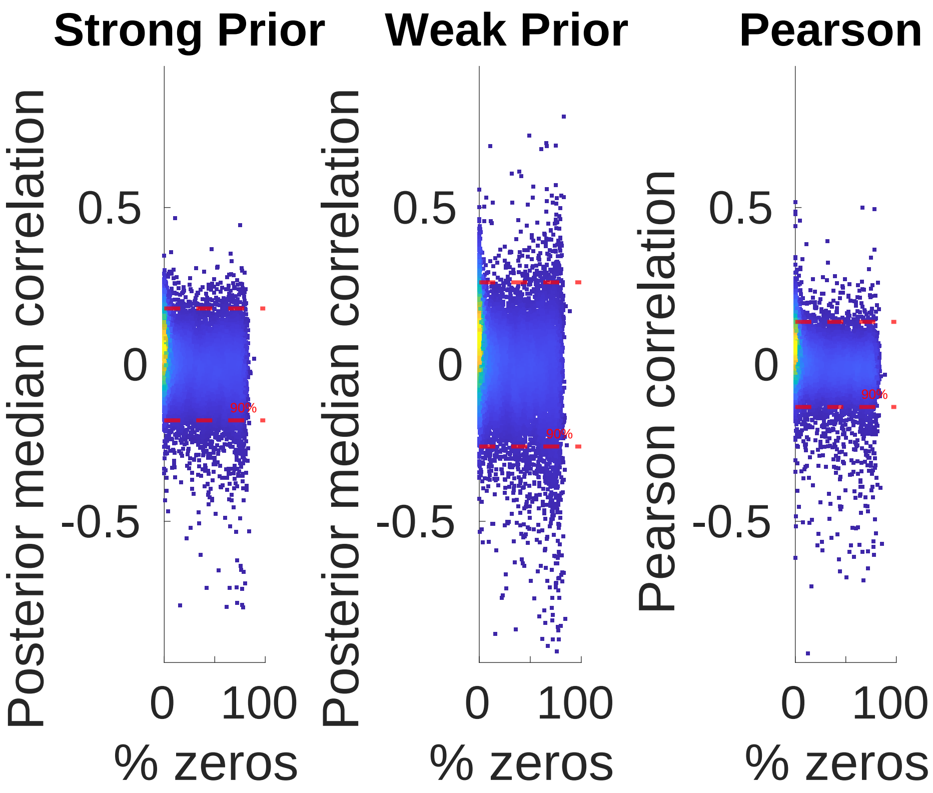


Figure S7: Posterior median correlation as function of % of zeros in raw data for strong prior (left), weak prior (middle). For comparison we also plot Pearson correlation as a function of % of zeros. Each dot represents a gene.

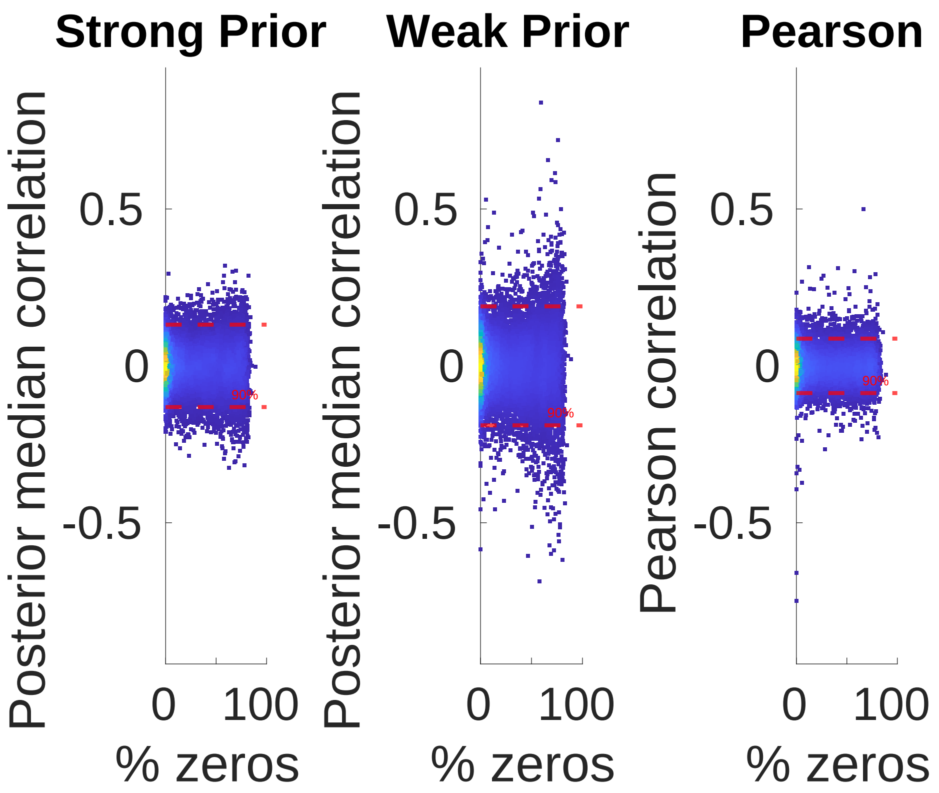


Figure S8: Posterior median correlation as function of % of zeros in raw negative control data for strong prior (left), weak prior (middle). For comparison we also plot Pearson correlation as a function of % of zeros. Each dot represents a gene.

### S6 Comparing predictions between alternative approaches

In the process of understanding SCRaPL's, we had to analyze different scenarios summarizing model's behavior. We consider the set of associations which are called as significant by at least one method, and split it into 3 categories: agreement between predictions, association labeling as significant by SCRaPL, but not Pearson, and vice-versa. Here we go through some important examples omitted from the main text, which demonstrate SCRaPL's superior performance.

#### S6.1 Classified as significant by SCRaPL and Pearson

In this section we present some cases where SCRaPL agrees with Pearson. For many of the examples we have good coverage for both methylation and expression. Furthermore we tend to have observations for a wide range of methylation and expression values.

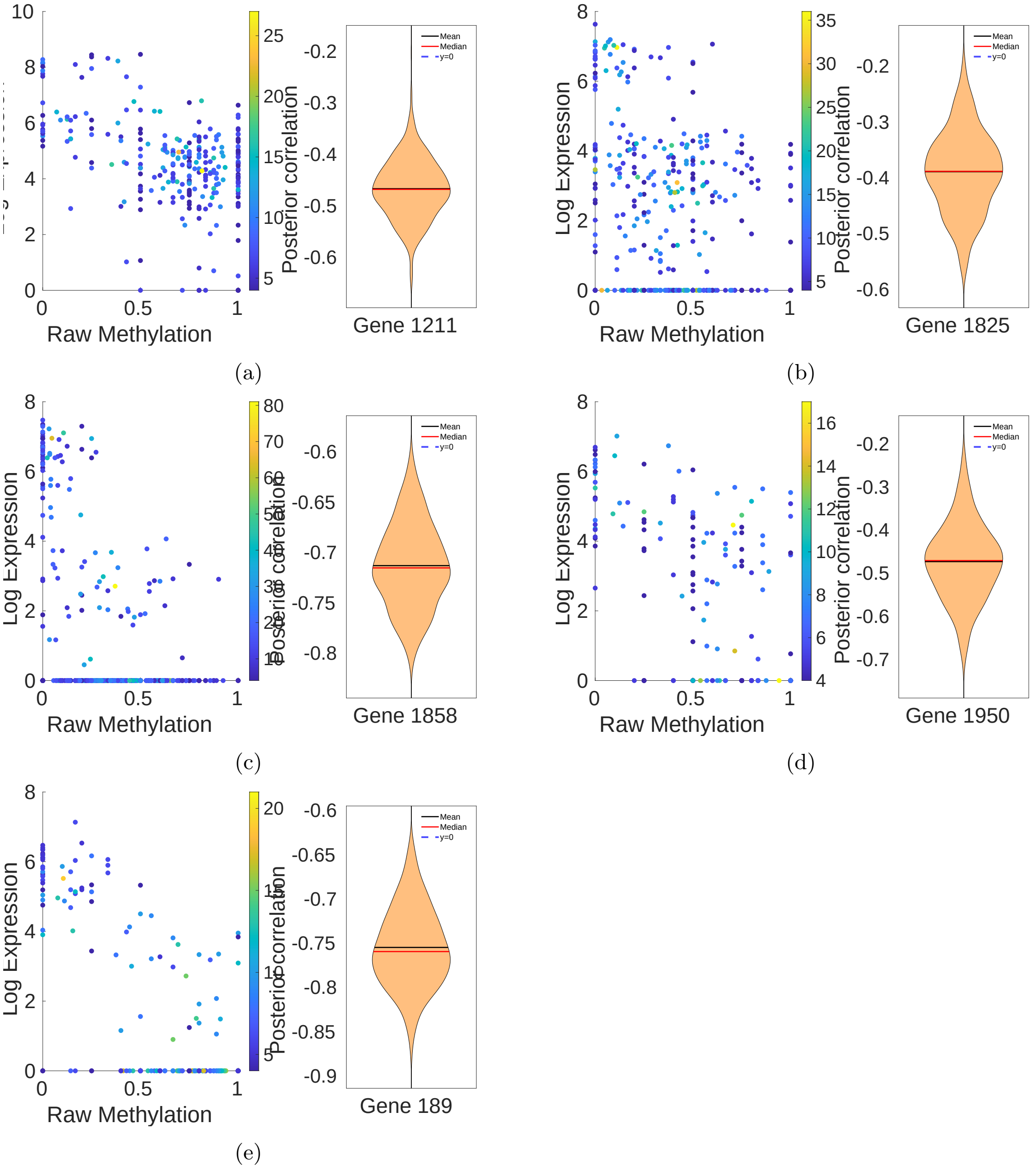


Figure S9: Examples where SCRaPL and Pearson agree on the significance of feature. In all figures the left part is a scatter plot of feature's raw data colorcoded by CpG coverage with normalized expression in the $log(1+x)$ scale. On the right we see the posterior for the same feature as infered by SCRaPL. Above figures include features with labels 1211 (S9a), 1825 (S9b), 1858 (S9c), 1950 (S9d) and 189 (S9e)

#### S6.2 Classified as insignificant by SCRaPL and significant by Pearson

In this subsection we generally have features with observations leading to spurious correlations. Some of these problems encountered in observations include, low expression/ number of observations, extremely low coverage and large number of zeros. For many of these features we see that are large areas where we do not have any observation for both methylation and expression.

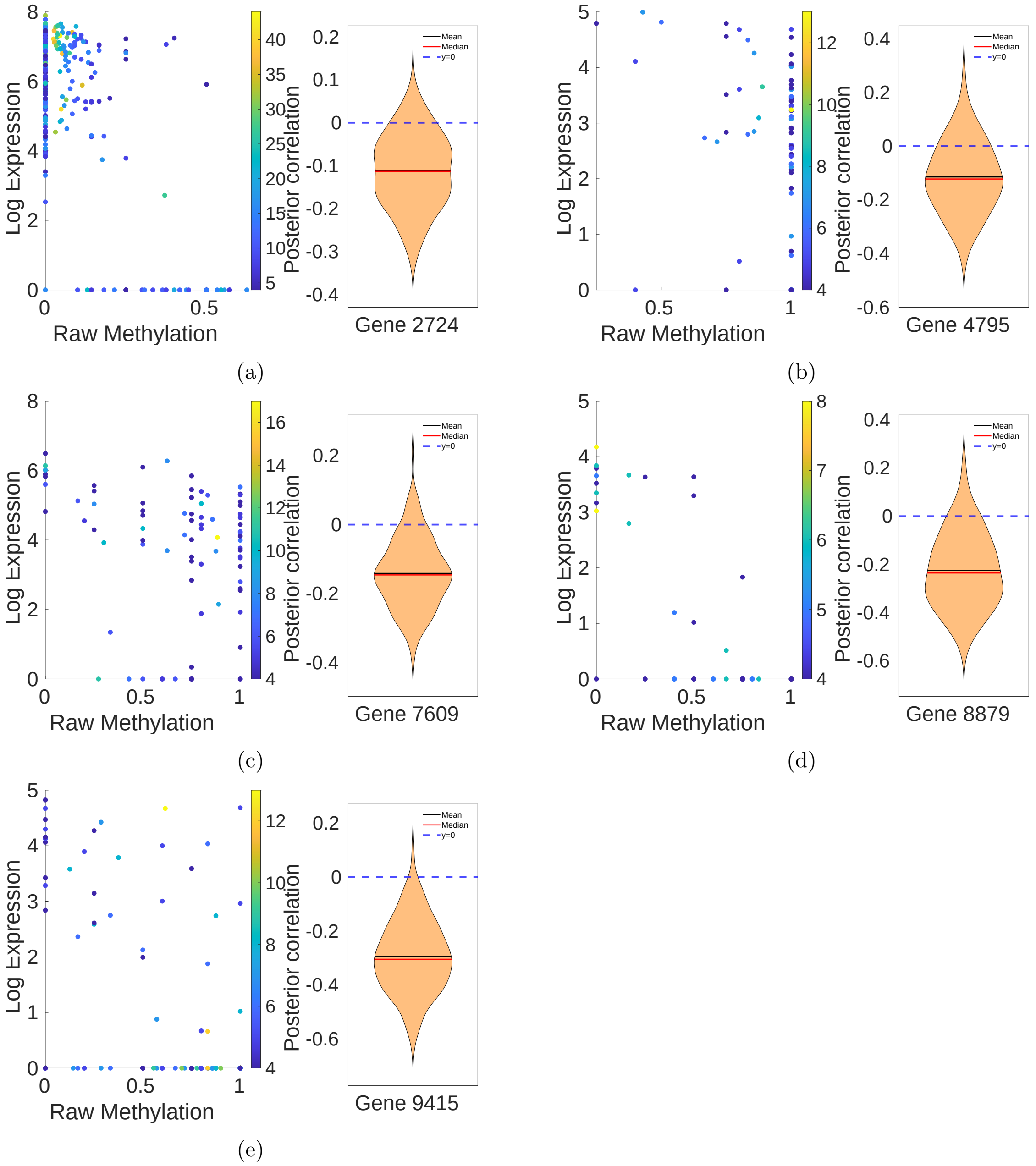


Figure S10: Examples which were classified as insignificant by SCRaPL and significant by Pearson correlation. In all figures the left part is a scatter plot of feature's raw data colorcoded by CpG coverage with normalized expression in the $log(1 + x)$ scale. On the right we see the posterior for the same feature as infered by SCRaPL. Above figures include features with labels 2724 (S10a), 4759 (S10b), 7609 (S10c), 8879 (S10d) and 9415 (S10e)

#### S6.3 Classified as significant by EFDR and insignificant by FDR

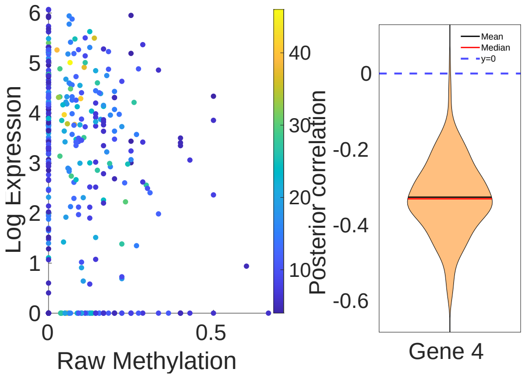


(a)

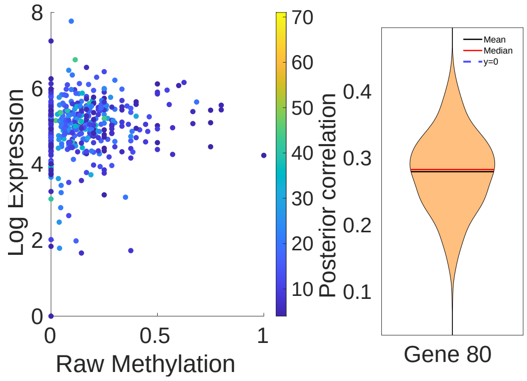


(b)

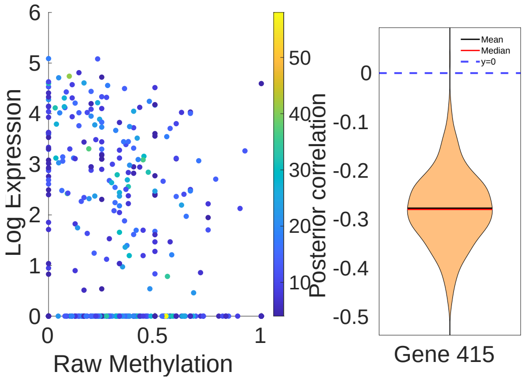


(c)

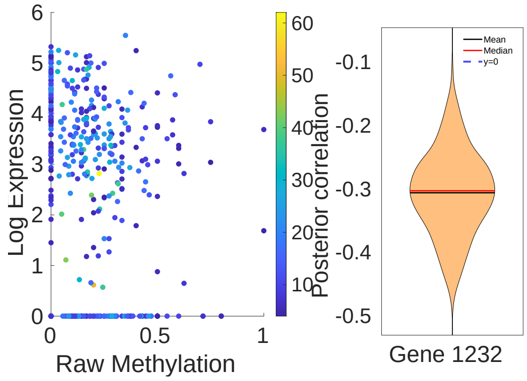


(d)

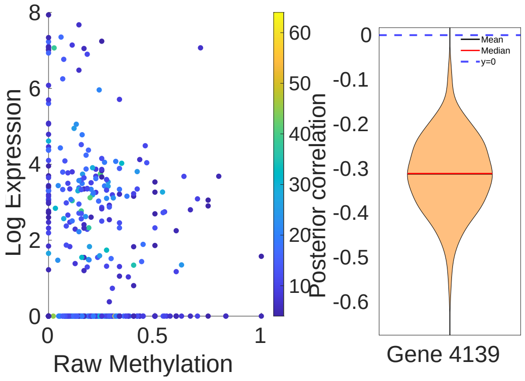


(e)

Figure S11: Examples which were classified as significant by SCRaPL and insignificant by Pearson correlation.In all figures the left part is a scatter plot of feature's raw data colorcoded by CpG coverage with normalized expression in the $log(1+x)$ scale. On the right we see the posterior for the same feature as infered by SCRaPL. Above figures include features with labels 4 (S11a), 80 (S11b), 415 (S11c), 1232 (S11d) and 4139 (S11e)

##### S6.4 Posterior correlation inference for common genes in bibliography

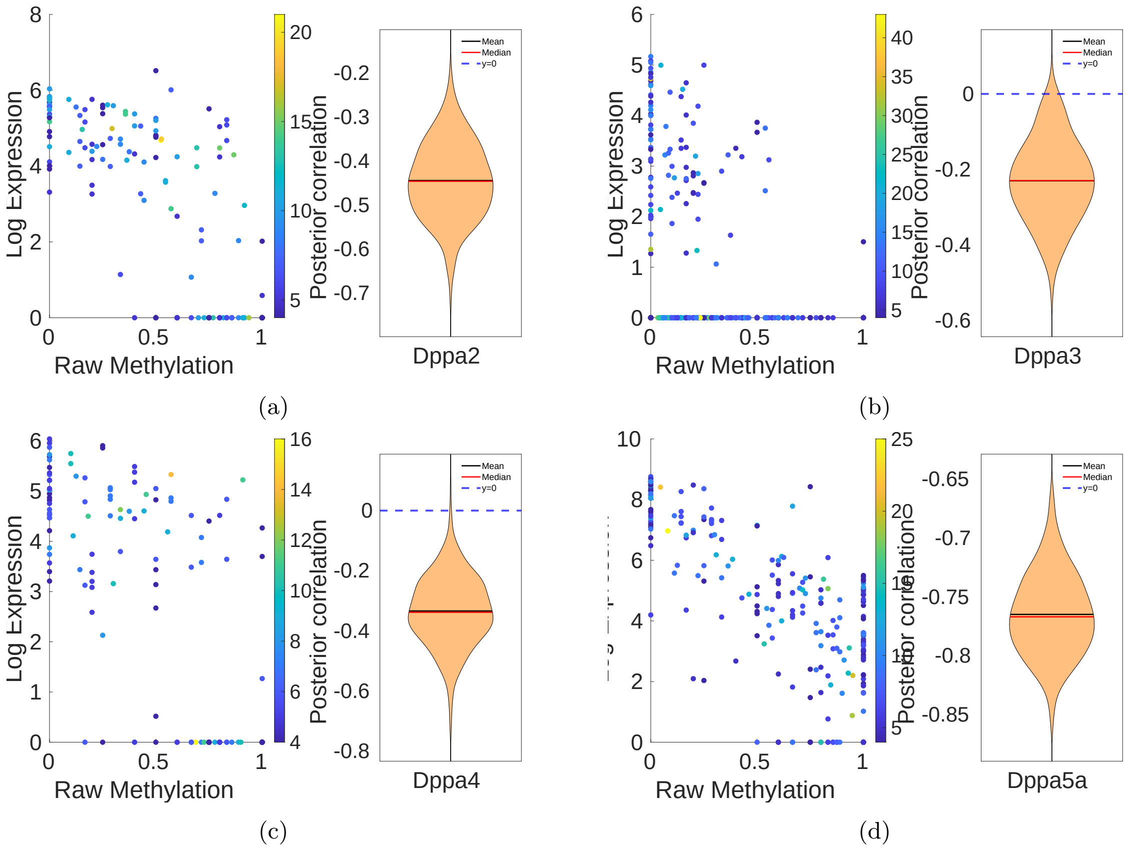


Figure S12: Raw data and posterior gene correlation for selected members of the Dppa gene family inferred with SCRaPL.

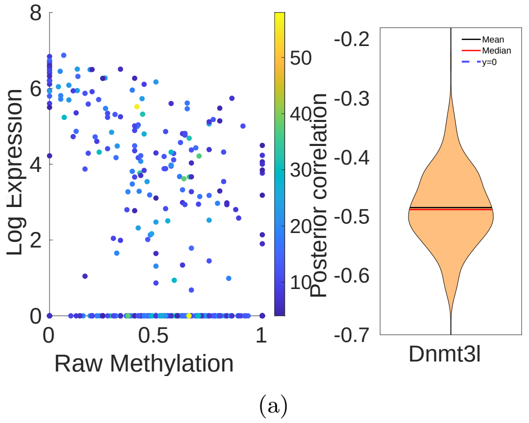


Figure S13: Raw data and posterior gene correlation for selcted members of the Dnmt gene family inferred with SCRaPL.

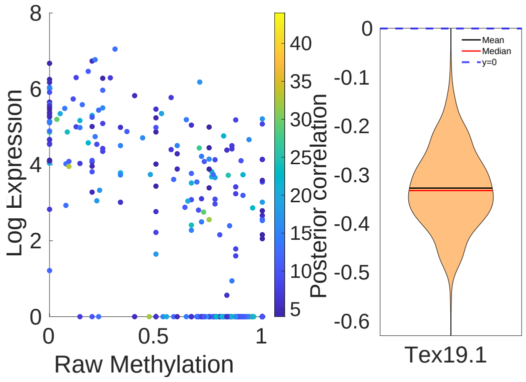


(a)

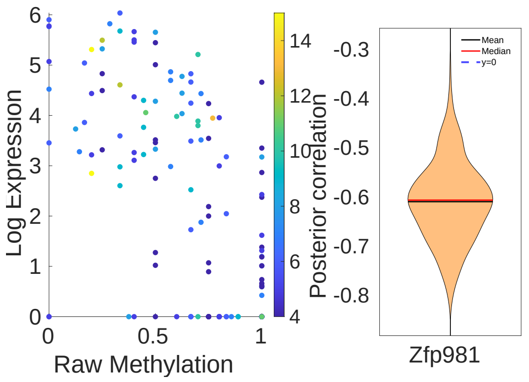


(b)

Figure S14: Raw data and posterior gene correlation for other studied genes Argelaguet et al. (2019) inferred with SCRaPL.

### S7 Gene Set Enrichment Analysis

To get a more spherical understanding of the genes with strong regulatory action identified by alternative approaches, we carry our Gene Set Enrichment Analysis (GSEA) analysis using DAVID Sherman et al. (2009). This process helps us to identify important biological processes by looking at over-represented genes in a starting pool of genetic markers. Using SCRaPL correlation outcomes we can determine a lists of biological outcomes that practitioners would discover. The $\gamma$ threshold for SCRaPL was set to 0.205 or 90% quantile of the folded distribution constructed by correlation samples of the permuted dataset.

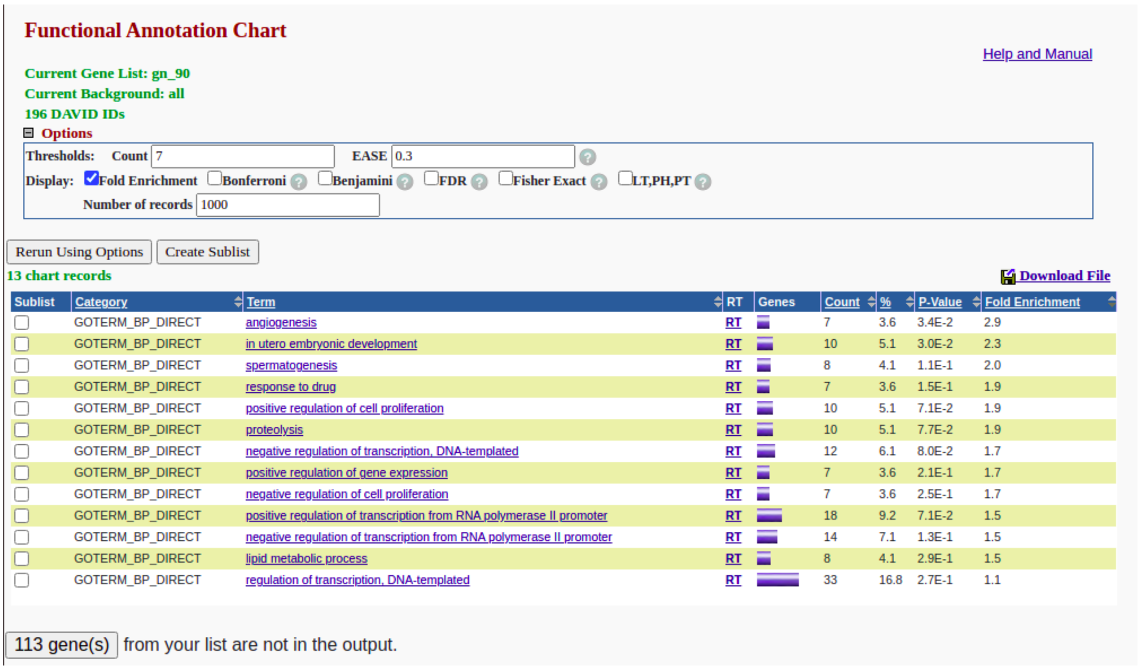

**Functional Annotation Chart**

Help and Manual

**Current Gene List: gn_90**
**Current Background: all**
**196 DAVID IDs**
**Options**

Thresholds: Count 7 EASE 0.3
Display: Fold Enrichment, Bonferroni, Benjamini, FDR, Fisher Exact, LT,PH,PT
Number of records 1000

Rerun Using Options | Create Sublist

**13 chart records**

Download File

| Sublist | Category | Term | RT | Genes | Count | % | P-Value | Fold Enrichment |
|---|---|---|---|---|---|---|---|---|
| | GOTERM_BP_DIRECT | angiogenesis | RT | | 7 | 3.6 | 3.4E-2 | 2.9 |
| | GOTERM_BP_DIRECT | in utero embryonic development | RT | | 10 | 5.1 | 3.0E-2 | 2.3 |
| | GOTERM_BP_DIRECT | spermatogenesis | RT | | 8 | 4.1 | 1.1E-1 | 2.0 |
| | GOTERM_BP_DIRECT | response to drug | RT | | 7 | 3.6 | 1.5E-1 | 1.9 |
| | GOTERM_BP_DIRECT | positive regulation of cell proliferation | RT | | 10 | 5.1 | 7.1E-2 | 1.9 |
| | GOTERM_BP_DIRECT | proteolysis | RT | | 10 | 5.1 | 7.7E-2 | 1.9 |
| | GOTERM_BP_DIRECT | negative regulation of transcription, DNA-templated | RT | | 12 | 6.1 | 8.0E-2 | 1.7 |
| | GOTERM_BP_DIRECT | positive regulation of gene expression | RT | | 7 | 3.6 | 2.1E-1 | 1.7 |
| | GOTERM_BP_DIRECT | negative regulation of cell proliferation | RT | | 7 | 3.6 | 2.5E-1 | 1.7 |
| | GOTERM_BP_DIRECT | positive regulation of transcription from RNA polymerase II promoter | RT | | 18 | 9.2 | 7.1E-2 | 1.5 |
| | GOTERM_BP_DIRECT | negative regulation of transcription from RNA polymerase II promoter | RT | | 14 | 7.1 | 1.3E-1 | 1.5 |
| | GOTERM_BP_DIRECT | lipid metabolic process | RT | | 8 | 4.1 | 2.9E-1 | 1.5 |
| | GOTERM_BP_DIRECT | regulation of transcription, DNA-templated | RT | | 33 | 16.8 | 2.7E-1 | 1.1 |

113 gene(s) from your list are not in the output.

(a)

Figure S15: GO analysis with features detected by SCRaPL.

The detected biological presented in figure S15 have at 7 or more genes associated with them and maximum p-value 0.3. Then processes were sorted based on their enrichment score (ie. how many times larger is the detected set compared to a random subset of the pool of genes linked to a particular biological process). SCRaPL has detected many processes directly linked to regulation of transcription and regulation of transcription linked to promoter regions (something expected as we are looking at promoters of a methylation-expression pair at early development where methylation plays a crucial role). Apart from them there are also biological processes linked to development, like in utero embryonic development and angiogenesis with high enrichment score. For the exact same filtering parameter and genes detected with Pearson correlation, the GO would indicated that "regulation of transcription, DNA-templated" is significant with enrichment score 1.5.

### S8 Connecting SCRaPL error model to likelihoods currently employed by practitioners

In the field of transcriptomics it is widely accepted that over-dispersion is an important feature of state of the art models Svensson (2020). Following the example of several papers He and Kulminski (2020); Svensson (2020); Vallejos et al. (2016) where they provide evidence that their model can handle excessive zeros by using over-dispersion, we demonstrate that zero-inflated Poisson is also a valid alternative as the over-dispersion property exists. For the sake of completeness we provide mean and variance formulas for the binomial posterior.

#### S8.1 Zero-inflated Poisson

$$\mathbb{P}(y) = \begin{cases} \int_{-\infty}^{\infty} \pi + (1-\pi)e^{-ae^x}\mathbb{N}\left(x,\mu,\sigma^2\right)dx, \textbf{if } y=0 \\ \int_{-\infty}^{\infty} \frac{1-\pi}{y!}a^y e^{yx-ae^x}\mathbb{N}\left(x,\mu,\sigma^2\right)dx, \textbf{else} \end{cases} \tag{1}$$

$$\begin{aligned}
\mathbb{E}(y) = \sum_{y=0}^{\infty} y\mathbb{P}(y) = \sum_{y=1}^{\infty} y\mathbb{P}(y) &= \sum_{y=1}^{\infty} y\frac{1-\pi}{y!}a^y \int_{-\infty}^{\infty} e^{yx-ae^x}\mathbb{N}\left(x,\mu,\sigma^2\right)dx \\
&= \sum_{y=1}^{\infty} \frac{1-\pi}{(y-1)!}a^y \int_{-\infty}^{\infty} e^{yx-ae^x}\mathbb{N}\left(x,\mu,\sigma^2\right)dx \\
&= a(1-\pi)\int_{-\infty}^{\infty} e^x e^{-ae^x}\mathbb{N}\left(x,\mu,\sigma^2\right)\sum_{y=1}^{\infty} a^{y-1}\frac{e^{(y-1)x}}{(y-1)!}dx \\
&= a(1-\pi)\int_{-\infty}^{\infty} e^x e^{-ae^x}\mathbb{N}\left(x,\mu,\sigma^2\right)\sum_{y=0}^{\infty} a^y\frac{e^{yx}}{y!}dx \\
&= a(1-\pi)\int_{-\infty}^{\infty} e^x e^{-ae^x}\mathbb{N}\left(x,\mu,\sigma^2\right)e^{ae^x}dx \\
&= a(1-\pi)\int_{-\infty}^{\infty} e^x\mathbb{N}\left(x,\mu,\sigma^2\right)dx = a(1-\pi)e^{\mu+\frac{\sigma^2}{2}}
\end{aligned}$$

Hence,

$$\mathbb{E}(y) = a(1-\pi)e^{\mu+\frac{\sigma^2}{2}} = m \tag{2}$$

$$\mathbb{E}\left(y^2\right) = \sum_{y=0}^{\infty} y^2\mathbb{P}(y) = \sum_{y=1}^{\infty} y^2\mathbb{P}(y)$$

$$= \sum_{y=1}^{\infty} y^2\frac{1-\pi}{y!}\int_{-\infty}^{\infty} a^y e^{yx-ae^x}\mathbb{N}\left(x,\mu,\sigma^2\right)dx = \sum_{y=1}^{\infty} y\frac{1-\pi}{(y-1)!}\int_{-\infty}^{\infty} a^y e^{yx-ae^x}\mathbb{N}\left(x,\mu,\sigma^2\right)dx$$

$$= a(1-\pi)\int_{-\infty}^{\infty} e^x e^{-ae^x}\mathbb{N}\left(x,\mu,\sigma^2\right)\sum_{y=0}^{\infty}\frac{(y+1)a^y e^{yx}}{y!}dx$$

$$= a(1-\pi)\int_{-\infty}^{\infty} e^x e^{-ae^x}\mathbb{N}\left(x,\mu,\sigma^2\right)\sum_{y=0}^{\infty}\frac{ya^y e^{yx}}{y!}dx + a(1-\pi)\int_{-\infty}^{\infty} e^x e^{-ae^x}\mathbb{N}\left(x,\mu,\sigma^2\right)\sum_{y=0}^{\infty}\frac{a^y e^{yx}}{y!}dx$$

$$= a(1-\pi)\int_{-\infty}^{\infty} e^x e^{-ae^x}\mathbb{N}\left(x,\mu,\sigma^2\right)\sum_{y=1}^{\infty}\frac{a^y e^{yx}}{(y-1)!}dx + a(1-\pi)\int_{-\infty}^{\infty} e^x\mathbb{N}\left(x,\mu,\sigma^2\right)dx$$

$$= a^2(1-\pi)\int_{-\infty}^{\infty} e^{2x}e^{-ae^x}\mathbb{N}\left(x,\mu,\sigma^2\right)\sum_{y=0}^{\infty}\frac{a^y e^{yx}}{y!}dx + a(1-\pi)\int_{-\infty}^{\infty} e^x\mathbb{N}\left(x,\mu,\sigma^2\right)dx$$

$$= a^2(1-\pi)\int_{-\infty}^{\infty} e^{2x}\mathbb{N}\left(x,\mu,\sigma^2\right)dx + a(1-\pi)\int_{-\infty}^{\infty} e^x\mathbb{N}\left(x,\mu,\sigma^2\right)dx = a(1-\pi)\left[ae^{2\mu+2\sigma^2} + e^{\mu+\frac{\sigma^2}{2}}\right]$$

$$\mathbb{V}(y) = \mathbb{E}\left(y^2\right) - \mathbb{E}(y)^2 = a(1-\pi)e^{\mu+\frac{\sigma^2}{2}}\left(1 + ae^{\mu+\frac{3\sigma^2}{2}}\right) - a^2(1-\pi)^2 e^{2\mu+\sigma^2}$$
$$= a(1-\pi)e^{\mu+\frac{\sigma^2}{2}}\left[1 + ae^{\mu+\frac{3\sigma^2}{2}} - a(1-\pi)e^{\mu+\frac{\sigma^2}{2}}\right]$$
$$= a(1-\pi)e^{\mu+\frac{\sigma^2}{2}}\left[1 + ae^{\mu+\frac{\sigma^2}{2}}\left(e^{\sigma^2} - 1 + \pi\right)\right] = m\left(1 + m\frac{e^{\sigma^2} - 1 + \pi}{1-\pi}\right)$$

Hence,

$$\mathbb{V}(y) = m\left(1 + m\frac{e^{\sigma^2} - 1 + \pi}{1-\pi}\right) \tag{3}$$

#### S8.2 Binomial

$$\mathbb{P}(k) = \int_{-\infty}^{\infty} \binom{n}{k} \Phi(x)^k \left(1 - \Phi(x)\right)^{n-k} \mathbb{N}\left(x, \mu, \sigma^2\right) dx \tag{4}$$

$$\mathbb{E}(k) = \sum_{k=0}^{n} k\mathbb{P}(k) = \int_{-\infty}^{\infty} \mathbb{N}\left(x, \mu, \sigma^2\right) \sum_{k=0}^{n} k\binom{n}{k} \Phi(x)^k \left(1 - \Phi(x)\right)^{n-k} dx$$
$$= n\int_{-\infty}^{\infty} \Phi(x)\mathbb{N}\left(x, \mu, \sigma^2\right) dx = n\Phi\left(\frac{\mu}{\sqrt{1+\sigma^2}}\right)$$

Hence,

$$\mathbb{E}(k) = n\Phi\left(\frac{\mu}{\sqrt{1+\sigma^2}}\right) \tag{5}$$

$$\mathbb{E}(k(k-1)) = \sum_{k=0}^{n} k(k-1)\mathbb{P}(k) = \int_{-\infty}^{\infty} \mathbb{N}\left(x, \mu, \sigma^2\right) \sum_{k=0}^{n} k(k-1)\binom{n}{k} \Phi(x)^k \left(1 - \Phi(x)\right)^{n-k} dx$$
$$= n(n-1)\int_{-\infty}^{\infty} \Phi(x)^2\mathbb{N}\left(x, \mu, \sigma^2\right) = n(n-1)\left[\Phi\left(\frac{\mu}{\sqrt{1+\sigma^2}}\right) - 2T\left(\frac{\mu}{\sqrt{1+\sigma^2}}, \frac{1}{\sqrt{1+2\sigma^2}}\right)\right],$$
$$T(h, a) = \mathbb{N}(h, 0, 1)\int_0^a \frac{\mathbb{N}(hx, 0, 1)}{1+x^2} dx$$

$$\mathbb{V}(k) = \mathbb{E}\left(k^2\right) - \mathbb{E}(k)^2 = \mathbb{E}(k(k-1)) + \mathbb{E}(k) - \mathbb{E}(k)^2$$
$$= n(n-1)\left[\Phi\left(\frac{\mu}{\sqrt{1+\sigma^2}}\right) - 2T\left(\frac{\mu}{\sqrt{1+\sigma^2}}, \frac{1}{\sqrt{1+2\sigma^2}}\right)\right] + n\Phi\left(\frac{\mu}{\sqrt{1+\sigma^2}}\right) - n^2\Phi\left(\frac{\mu}{\sqrt{1+\sigma^2}}\right)^2$$
$$= n^2\Phi\left(\frac{\mu}{\sqrt{1+\sigma^2}}\right)\left[1 - \Phi\left(\frac{\mu}{\sqrt{1+\sigma^2}}\right)\right] - 2n(n-1)T\left(\frac{\mu}{\sqrt{1+\sigma^2}}, \frac{1}{\sqrt{1+2\sigma^2}}\right)$$

hence,

$$\mathbb{V}(k) = n^2\Phi\left(\frac{\mu}{\sqrt{1+\sigma^2}}\right)\left[1 - \Phi\left(\frac{\mu}{\sqrt{1+\sigma^2}}\right)\right] - 2n(n-1)T\left(\frac{\mu}{\sqrt{1+\sigma^2}}, \frac{1}{\sqrt{1+2\sigma^2}}\right)$$

### S9 Null hypothesis testing

#### S9.1 Pearson

In most settings where Pearson correlation with hypothesis testing is applied, the aim is to determine whether the estimated value of correlation is generated from an uncorrelated bivariate normal distribution. Here we are interested to investigate the more complicated null hypothesis that the data generating correlation lives in an interval around 0. Hence the first step is to determine the distribution of sampled Pearson correlations $r$ given true correlation $\rho$ in a correlated bivariate normal distribution. According to Hotelling (1953) that distribution is:

$$f(r, \rho) = \frac{(n-2)\Gamma(n-1)(1-\rho^2)^{\frac{n-1}{2}}(1-r^2)^{\frac{n-2}{4}}}{\sqrt{2\pi}\Gamma(n-\frac{1}{2})(1-\rho r)^{n-\frac{3}{2}}} {}_2F_1\left(\frac{1}{2}, \frac{1}{2}, \frac{2n-1}{2}, \frac{1+r\rho}{2},\right) \tag{6}$$

where $\Gamma$ is the gamma function and ${}_2F_1$ is the Gaussian hypergeometric function. In the special case of $\rho = 0$ we get student a reparametrized version of student t-distribution. However we are interested in the null distribution of correlations with magnitude below a threshold $\gamma_{prs}$. The get it, we integrate $f(r, \rho)$ over that range $[-\gamma_{prs}, \gamma_{prs}]$.

$$p(r) = \frac{1}{\mathbb{Z}} \int_{-\gamma_{prs}}^{\gamma_{prs}} f(r, \rho) d\rho \tag{7}$$

Where $\mathbb{Z} = \int_{-1}^{1} p(r)dr$. Integrating $f(r, \rho)$ over $\rho$ is not trivial as there is not closed form solution. Hence we resort to numerical integration using Matlab's built in function. The p-value under the null hypothesis for a Pearson correlation estimate $p_{cor}$ is

$$\mathbb{P}(R \geq |p_{cor}|) = \int_{-1}^{-|p_{cor}|} p(r)dr + \int_{|p_{cor}|}^{1} p(r)dr = 2 \int_{-1}^{-|p_{cor}|} p(r)dr \tag{8}$$

To simplify this integral and get the simplified expression of the right hand side, we use that $f(r, \rho) = f(-r, -\rho)$. Type I error is controlled with standard FDR Benjamini and Hochberg (1995) as in the case of $\gamma_{prs} = 0$.

##### S9.2 SCRaPL

The aim of SCRaPL is to identify genes with strong correlation across molecular layers using gene specific posterior correlation. Mathematically this is done by estimating the probability of correlation's magnitude (ie. $|\rho_j|$) being above a threshold $\gamma$.

$$p_j(\gamma) = \mathbb{P}(|\rho_j| \geq \gamma) \tag{9}$$

If $p_j(\gamma)$ is larger than a threshold $a$ then correlation on genomic region $j$ is labeled statistically significant. To estimate $\gamma$ we have a data driven approach in place. More precisely, we look various quantiles in negative control data. Using $\gamma$ we calibrate $\alpha$ such that EFDR Newton et al. (2004) is below 10%. Since $p_j(\gamma)$ is a cumulative density function, under the null hypothesis it is uniformly distributed. Hence we apply the same procedure in the original and negative control data and compare detection rates. This test becomes problematic in case $\gamma = 0$ as $p_j(\gamma) = 1$ for every $j$. In this case we apply the rule from Bochkina and Richardson (2007) based on the maximum posterior probabilities associated to the one-sided hypothesis $\rho_j > 0$ and $\rho_j < 0$, mathematically summarized as follows:

$$2\max(\pi_j, 1 - \pi_j) - 1 > a, \ \textbf{with} \ \pi_j = \mathbb{P}(\rho_j \geq 0) \tag{10}$$

Parameter $a$ is calibrated such that EFDR is below 10%. The max-rule in equation 10 is uniformly distributed under the null hypothesis. This limitation of this rule is that results are correct for the case of symmetric around 0 posterior correlation distributions. Therefore, we use it here as an approximation.

Liang He and Alexander Kulminski. Nebula: a fast negative binomial mixed model for differential expression and co-expression analyses of large-scale multi subject single-cell data. *bioRxiv*, 2020.

Harold Hotelling. New light on the correlation coefficient and its transforms. *Journal of the Royal Statistical Society. Series B (Methodological)*, 15(2):193–232, 1953.

Romain Lopez, Jeffrey Regier, Michael B Cole, Michael I Jordan, and Nir Yosef. Deep generative modeling for single-cell transcriptomics. *Nature methods*, 15(12):1053–1058, 2018.

Michael A Newton, Amine Noueiry, Deepayan Sarkar, and Paul Ahlquist. Detecting differential gene expression with a semiparametric hierarchical mixture method. *Biostatistics*, 5(2):155–176, 2004.

Sebastian Pott. Simultaneous measurement of chromatin accessibility, dna methylation, and nucleosome phasing in single cells. *Elife*, 6:e23203, 2017.

Brad T Sherman, Richard A Lempicki, et al. Systematic and integrative analysis of large gene lists using david bioinformatics resources. *Nature protocols*, 4(1):44, 2009.

Valentine Svensson. Droplet scrna-seq is not zero-inflated. *Nature Biotechnology*, 38(2):147–150, 2020.

Catalina A Vallejos, Sylvia Richardson, and John C Marioni. Beyond comparisons of means: understanding changes in gene expression at the single-cell level. *Genome biology*, 17(1):1–14, 2016.
